# Cooperative genetic networks drive a mammalian cell state transition

**DOI:** 10.1101/2020.03.23.000109

**Authors:** Andreas Lackner, Robert Sehlke, Marius Garmhausen, Giuliano Giuseppe Stirparo, Michelle Huth, Fabian Titz-Teixeira, Petra van der Lelij, Julia Ramesmayer, Henry Fabian Thomas, Meryem Ralser, Laura Santini, Mihail Sarov, A. Francis Stewart, Austin Smith, Andreas Beyer, Martin Leeb

**Author notes:** Equal contribution (Andreas Lackner, Robert Sehlke, Marius Garmhausen, Giuliano Giuseppe Stirparo). Equal contribution (Michelle Huth, Fabian Titz-Teixeira).

## Abstract

In the mammalian embryo, epiblast cells must exit their naïve state and acquire formative pluripotency. This cell state transition is recapitulated by mouse embryonic stem cells (ESCs), which undergo pluripotency progression in defined conditions *in vitro*. However, our understanding of the molecular cascades and gene-networks involved in the exit from naïve pluripotency remains fragmented. Here we employed a combination of genetic screens in haploid ESCs, CRISPR/Cas9 gene disruption, large-scale transcriptomics and computational systems-biology to delineate the regulatory circuits governing naïve state exit. Transcriptome profiles for 73 knockout ESC lines predominantly manifest delays on the trajectory from naive to formative epiblast. We find that gene networks operative in ESCs are active during transition from pre- to post-implantation epiblast *in utero*. We identified 374 naïve-associated genes tightly connected to epiblast state and largely conserved in human ESCs and primate embryos. Integrated analysis of mutant transcriptomes revealed funneling of multiple gene activities into discrete regulatory modules. Finally, we delineate how intersections with signaling pathways direct this pivotal mammalian cell state transition.

## Introduction

Mouse embryonic stem cells (ESCs) can self-renew in defined conditions in a ground state of naïve pluripotency ^1^. The ESC exit from naïve pluripotency provides an amenable experimental system for dissection of a cell fate decision paradigm ^2, 3^. Naïve pluripotency is under control of a gene regulatory network (GRN) containing the core pluripotency transcription factors (TFs) *Pou5f1*, *Sox2* and naïve-specific TFs like *Nanog*, *Esrrb*, *Klf4* and others ^4–6^. In defined cell culture conditions that include inhibitors against Mek1/2 (PD0325901) and Gsk3 (CHIR990201, CH; collectively termed ‘2i’), ESCs can be homogenously maintained in the naïve state ^7^. Within 24 to 36 hours after withdrawal of 2i, ESCs transit into formative pluripotency, entirely losing naïve identity ^2^. During this transition, the naïve GRN is extinguished and expression of formative factors like *Otx2*, *Pou3f1*, *Dnmt3a/b* and *Fgf5* is initiated. A similar transition is evident during peri-implantation development, where the TF-network maintaining naïve pluripotency dissolves between embryonic day (E) 4.5 and E5.5 ^8–,10^.

The speed of the naïve to formative GRN transition is notable because: i) the cell cycle is around 12 hours long; ii) all factors that are required to establish and maintain naïve pluripotency are expressed robustly in naïve cells; and (iii) the naïve pluripotency network is recursively self-reinforcing. The rapid dissolution of naïve pluripotency implies the existence of circuit-breaking mechanisms. In recent years, we and others have identified various factors promoting ESC differentiation using screens in haploid and diploid ES cells ^11–15^.

Robust assays employing ESCs expressing a Rex1-promoter driven destabilised GFP reporter ESC line (Rex1::GFPd2) enable the dissection of the exit from naïve pluripotency in high resolution ^2, 16^. Rex1-GFP downregulation is initiated within 24h after 2i withdrawal (N24) and completed after 48h (N48). Nevertheless, the exact nature, mechanistic underpinnings and sequence of events during exit from naïve pluripotency remain only partially understood. In particular, we lack insight into how the different molecular components of the system co-operate to elicit proper cell fate transition.

Here, we have driven a Rex1-GFP reporter screen to saturation, thus providing an extensive list of genes and pathways involved in the exit from naïve pluripotency. We utilized this information in a systems biology approach to explore regulatory principles of the exit from naïve pluripotency. To evaluate dependencies and causal relationships within the pluripotency and differentiation circuitries, we probed the response of the differentiation program to a comprehensive series of exit factor gene knockouts. Through computational integration of molecular profiling data with regulatory networks and *in vivo* GRN-trajectories, we expose the regulatory foundations of a cell fate choice paradigm at a pivotal junction in early mammalian development.

## Results

### Haploid ES cell saturation screen

Haploid ES cells are an efficient platform for insertional mutagenesis-based screens ^11, 17, 18^. We previously reported a medium-scale screen comprising approximately 5×10^4^ mutagenic events to identify factors regulating the exit from naïve pluripotency ^12^. We have now driven this approach to saturation by assaying approximately 1.2 million mutations in receptive genomic regions that cause delays in Rex1 downregulation in two independent Rex1-reporter cell lines (Figure 1ab), utilizing three different mutagenic vectors in 35 independent screens (Figure 1cd).

Stringent filtering resulted in a candidate list comprising 489 genes (Supplementary Table 1). Reassuringly, the known exit from naïve pluripotency regulators *Tcf7l1, Fgfr2, Jarid2* and *Mapk1* (*Erk2*) ^19^ were among the highest ranked (Supplementary Figure 1a). The candidate hit genes were enriched for processes involved in transcription regulation, epigenetics and signalling-related functions (Supplementary Figures 1bc; Supplementary Table 1), as well as RNA binding functions in line with emerging evidence of RNA regulatory mechanisms in cell fate control ^20^.

These screens generated a candidate inventory of the machinery that mediates exit from naïve pluripotency. Many identified genes are not specific to pluripotency and have functions in common pathways and processes, implying that the exit from naïve pluripotency utilizes widely expressed cellular machinery. Therefore, mechanisms mediating ESC transition might also be utilised in other differentiation processes.

### Establishment of a mutant ESC library for systematic transcriptional profiling

To characterise deficiencies in naïve exit, we generated KO ESC lines deficient for 73 selected genes, comprising top ranked genes from the mutagenic screen together with components from pathways and protein complexes for which multiple members were recovered, even if just below the cut-off threshold (e.g. the Paf complex member *Leo1*, the mTORC1 regulator *Tsc2* and the NMD component *Smg6*), and *Mbd3*, *Zfp281* and *L3mbtl3* as known players in the exit from naïve pluripotency ^14^. Three control genes were included that are either not expressed in ES cells (Nestin), expected to be neutral (Hprt), or whose ablation was expected to accelerate differentiation (cMyc). Paired gRNAs were used to disrupt target genes in a diploid <u>R</u>ex1::GFPd2 reporter ES cell line carrying a <u>C</u>as<u>9</u>-transgene (henceforth termed RC9 cells) (Figure 1a). Following an efficient parallelized approach, we established passage matched and isogenic cell lines, thus maximising comparability (Supplementary Figure 1de).

**Figure 1.**
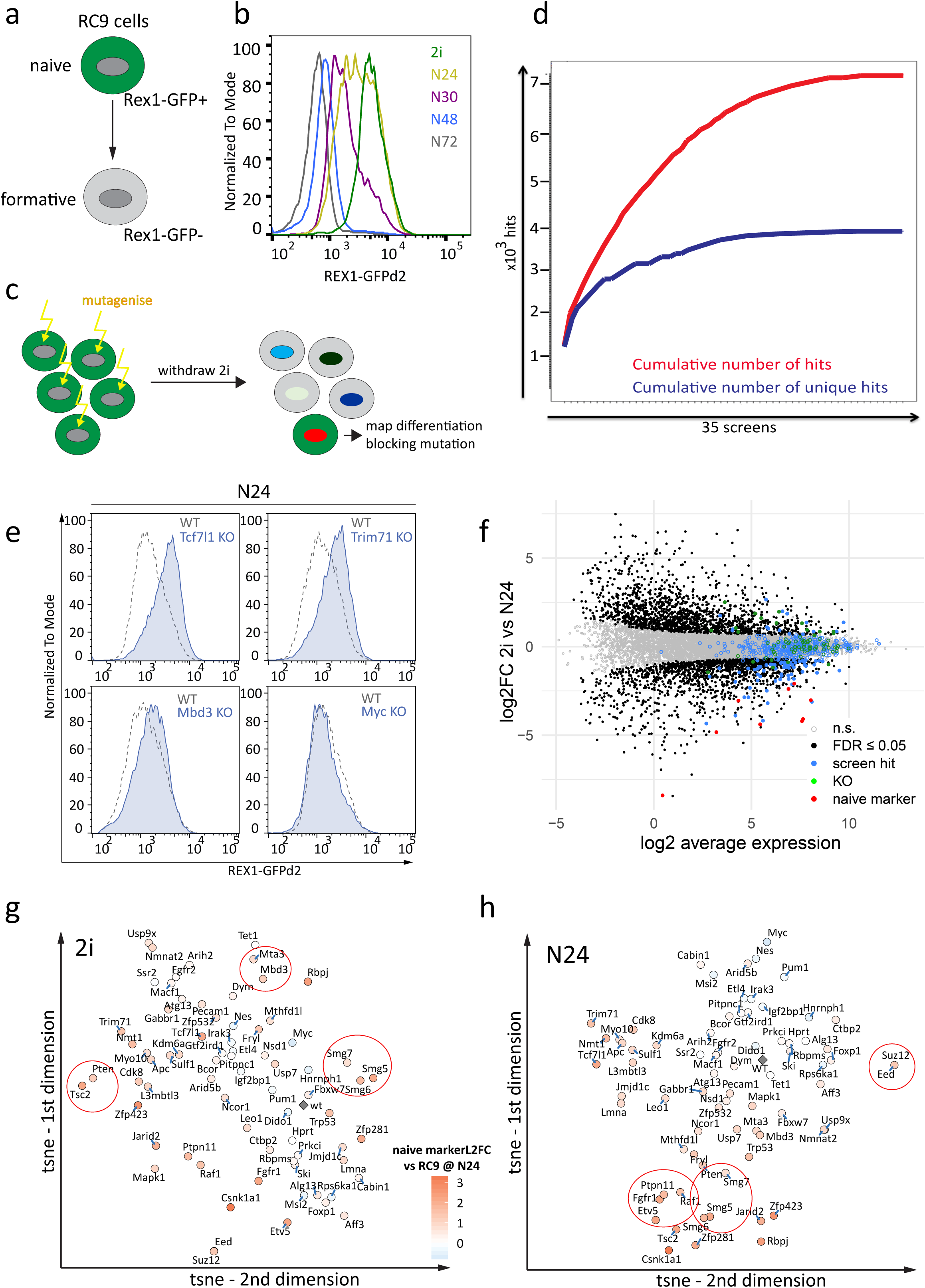
**a,** Illustration of the Rex1::GFPd2 reporter cell line and its’ exit from naïve pluripotency. Rex1-GFPd2 (in short Rex1-GFP) expression is tightly linked to naïve pluripotency. Shutdown of GFP expression indicates commitment to differentiation. **b,** FACS analysis of Rex1-GFPd2 reporter levels throughout a 72h differentiation time-course after 2i withdrawal. **c,** Scheme of the screening strategy to identify candidate genes involved in exit from naïve pluripotency. After random insertional mutagenesis using piggyBac transposon gene-traps, haploid Rex1::GFPd2 ESCs were released into differentiation. Cells that maintain GFP expression after exposure to differentiation conditions were isolated and the gene-trap insertion sites mapped. **d,** Plot showing cumulative number of hits and the cumulative number of novel hits over 35 independent insertional mutagenesis screens in haploid ESCs. **e,** Representative Rex1-GFP FACS plots showing the differentiation delays 30h after 2i withdrawal of *Tcf7l1*, *Trim71* and *Mbd3* KOs. A Myc KO served as a negative control. Blue indicates the KO FACS profiles and dashed lines indicates WT Rex-GFP. **f,** Differential expression of genes at N24 vs. 2i in WT RC9 cells. Black dots show significance (FDR ≤ 0.05, H0: |FC| > 1.5). Pluripotency genes are red dots, haploid screen hits are blue dots, the 73 KO-genes are green dots. **g,** t-SNE projection of the 73 KOs in 2i, based on expression of 3058 differentiation-associated genes (RC9^N24^ vs. RC9^2i^, FDR ≤ 0.05, H0: |FC| > 1.5). Differentiation defects observed at N24 in the respective KOs are indicated by a colour gradient, measured as average naive marker log2fold change (based on expression levels of *Esrrb, Nanog, Tfcp2l1, Tbx3, Prdm14* and *Klf4*) in the respective KO at N24. Red: delayed differentiation; blue: accelerated differentiation - see key in **h,**. **h,** Similar to **g,** for KOs at N24

Full protein deficiency was validated for 14 KOs (*Eed, Suz12, Jarid2, Kdm6a, Smg5, Smg6, Smg7, Tsc2, Pten, Raf1, Tcf7l1, Leo1, Nmt1,* and *Csnk1a1*), while reduced protein expression in concordance with a heterozygote phenotype was confirmed for *Mapk1* (Supplementary Figure 1f). For five further genes (*Alg13, Dido1, Msi2, Etv5, Jmjd1c*) we confirmed absence of the corresponding transcripts or specific out of frame deletion of an exon by RT-qPCR or Sanger sequencing of RT-PCR products. Successful rescue experiments using 3xflag-tagged transgenes for six genes (*Rbpj, Etv5, Fgfr1, Jarid2, Mbd3,* and *Tcf7l1*) established causality between the observed genotype and phenotype (Supplementary Figures 1gh). Thus, all tested knockouts showed the expected impact on RNA or protein expression. However, we cannot exclude the possibility of hypomorphic phenotypes in some cases.

ESC differentiation behaviour is highly dependent on cell density and timing of medium changes. To enable robust comparison of the differentiation of multiple KO ESC lines in parallel, we performed differentiations batch wise in duplicates, always including WT ESCs and negative controls across seven experiments. At N24 we assayed the differentiation status by FACS analysis (Figure 1e) and extracted RNA for transcriptome analysis.

### The exit machinery is already poised in 2i

Batch corrected RNA-seq data comprising 14 replicates of WT ES cells indicated 3058 differentially expressed genes (DEG) between 2i and N24 (H0: |FC| < 1.5, FDR ≤ 0.05; Figure 1f, Supplementary Figure 1i,j and Supplementary Table 1). Interestingly, most of the 489 genes identified in the haploid screens including the 73 genes selected for KO did not change significantly in transcript expression between 2i and N24 and were not present in the list of DEG (Figures 1f and S1K), with only *Jarid2*, *L3mbtl3*, *Pten* and *Zfp423* showing upregulation at N24 (FC ≥ 2; adj.p. ≤ 0.05). This implies that the exit machinery is already embedded in the ground state and ESCs are poised for rapid decommissioning of naïve identity and entry into differentiation ^19^.

To facilitate interrogation of the KO gene expression datasets, we developed an interactive online tool (GENEES - <u>Ge</u>netic <u>N</u>etwork <u>E</u>xplorer for <u>E</u>mbryonic <u>S</u>tem Cells – http://shiny.cecad.uni-koeln.de:3838/stemcell_network/; User: stemcells; Pw: S3m7KmIfHr9).

Using t-Distributed Stochastic Neighbour Embedding (t-SNE), we visualized similarities between KOs based on expression of the DEG in 2i and at N24. We observed clustering of members of the same complex or pathway: *Eed-Suz12* (PRC2), *Ptpn11*-*Raf1*-*Fgfr1*-*Etv5* (Fgf/ERK), *Smg5*-*Smg6*-*Smg7* (NMD; nonsense mediated decay), *Mta3*-*Mbd3* (NuRD), *Pten*- *Tsc2* (mTorc1 signaling) (Figure 1gh). The transcription profiles of the KO ESCs clustered by culture condition (2i and N24) and to a lesser extent by genotype (Supplementary Figure 1i). This is consistent with the phenotypic observation that despite manifesting differentiation delays at N24, all of the KO ESCs ultimately departed from the naïve state during longer differentiation timecourses, as measured by loss of Rex1GFP. Furthermore, even KOs that showed extensive Rex1GFP downregulation delays at N24 displayed transcription profiles that were globally adjusted towards differentiation. Therefore, the knockout of a single gene is not sufficient to permanently block exit from naïve pluripotency in culture, in accord with the finding that ternary depletion of *Tcf7l1*, *Etv5* and *Rbpj* is required for sustained self-renewal in the absence of 2i or LIF ^21^.

### The exceptional role of *Csnk1a1* and the involvement of compensatory mechanisms

The *Csnk1a1* KO is an exception to this generalization, clustering with 2i samples at N24 (Supplementary Figure 1i). Csnk1a1 is a serine threonine kinase and a component of the beta-catenin destruction complex. Although KO of another destruction complex member *Apc*, or of the downstream repressor *Tcf7l1* resulted in the upregulation of similar gene-sets (Supplementary Figure 1l), we observed stronger differentiation defects and larger amplitude of gene deregulation in two independently derived *Csnk1a1* mutants. However, these mutants also exhibited markedly reduced population doubling rates, which was not shown by *Apc* or *Tcf7l1* mutants. Upon continuous culture in 2i medium (∼5 passages), proliferation was restored in *Csnk1a1* KO cells and differentiation potential was regained, suggesting up-regulation of compensatory mechanisms and a likely effect of proliferation rate on differentiation kinetics. Csnk1a1 can be chemically inactivated using Epiblastin A, ^22^. We found that Epiblastin A partially blocked exit from naïve pluripotency, without affecting proliferation within the duration of the assay (Supplementary Figure 1m). A second case of phenotype adaptation was observed with Pum1. *Pum1* KOs showed pronounced differentiation delays during early passages (Supplementary Figure 1n), as also seen for acute Pum1 depletion by siRNA and in previously generated CRISPR KO ESCs ^12^. However, the phenotype was lost in later passages and *Pum1* KO cells showed WT like expression-levels of naïve marker genes (Figures 2a and S1a).

**Figure 2.**
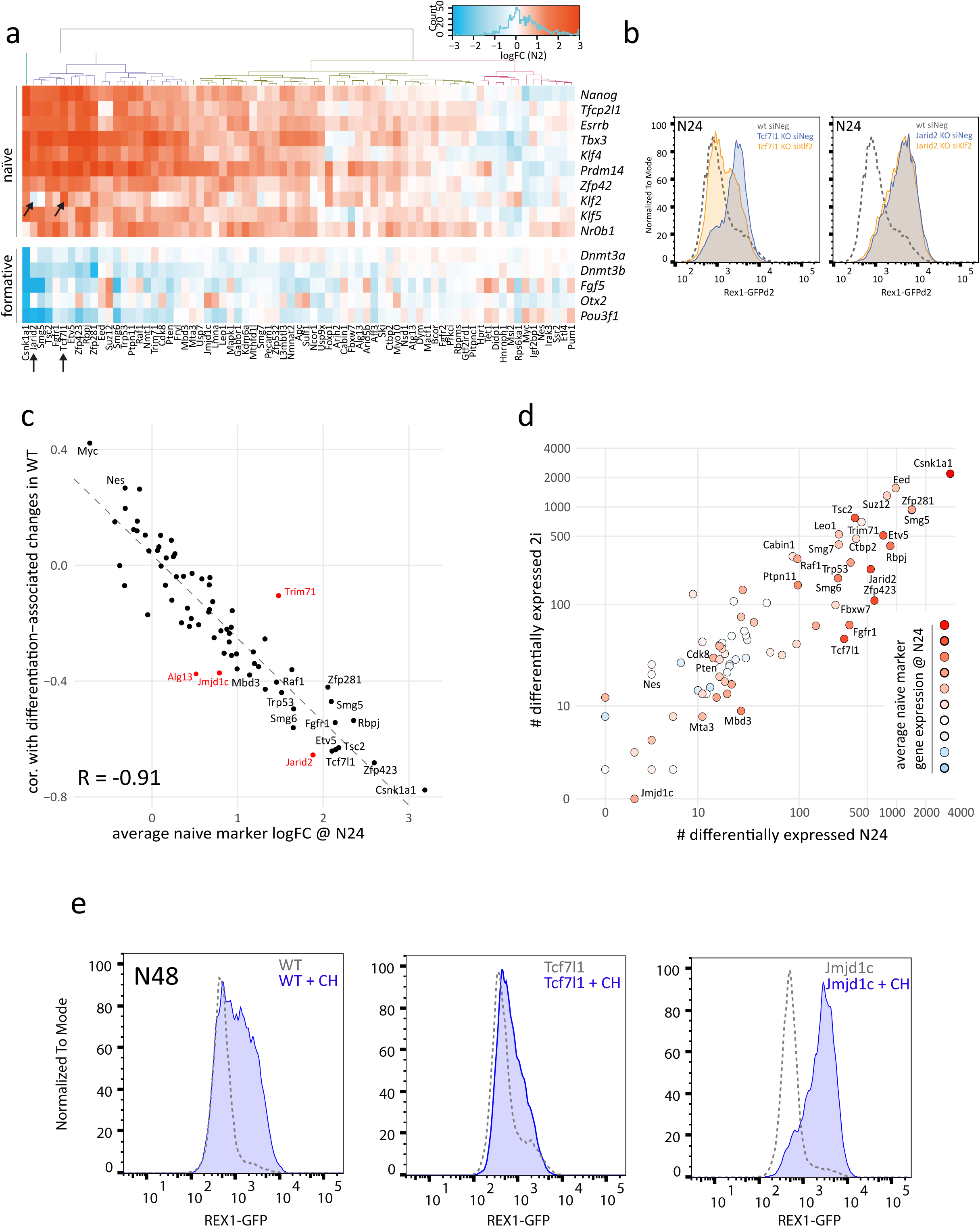
**a,** Naïve (top) and formative (bottom) marker gene expression changes at N24 compared to WT in all 73 KO ESCs. Clustering based on naïve marker gene expression shows a wide distribution of differentiation delay phenotypes. **b,** Rex1GFP FACS analysis of *Jarid2* and *Tcf7l1* KO ESCs after transfection with negative control or *Klf2* specific siRNAs. **c,** For each of the 73 KOs, the change of mean naïve marker gene expression at N24 (KO vs WT) was plotted against the change of the global transcriptome at N24 state (correlation of KO-induced change at N24 with differentiation-induced changes in WT differentiation, N24 vs 2i). **d,** Comparison of number of genes deregulated in 2i and at N24 in all KOs (FDR ≤ 0.05, H0: |FC| < 1.5). Differentiation phenotypes are colour coded according to mean naïve marker gene expression change in KOs at N24. **e,** Rex1GFP FACS analysis of WT, *Jmjd1c* and *Tcf7l1* KO cells at N48 cultured with and without the Gsk3 inhibitor, CH.

### Robust feedback wiring in the naïve TF network

Interestingly, the transcriptome data revealed that exit factors do not, in general, reduce naïve transcription factor (TF) expression in the ground state (Supplementary Figure 2a). *Rbpj* and *Trim71* KO resulted in moderate increases in *Klf4*, while the aforementioned *Csnk1a1* KO ESCs showed limited up-regulation of both *Klf4* and *Tbx3* in 2i. Other KOs had only minor effects on the naïve network in 2i. These data are consistent with robust feedback wiring in the naïve TF network ^4, 5^ and neutralisation of most ‘differentiation drivers’ in 2i in culture ^23^. In contrast, we observed a more extensive impact on formative markers. Several KO cell lines showed lower baseline expression in 2i of *Otx2, Fgf5, Dnmt3a/b* and *Pou3f1* (Oct6) (Supplementary Figure 2b). In line with recent results, we noted that depletion of several Fgf/ERK components resulted in reduced *Dnmt3a/b, Pou3f1* and *Fgf5* expression in 2i ^21^. Although Fgf/ERK signalling is effectively inhibited in 2i ^7^, our data suggest that either residual pathway activity or potential moonlighting functions of pathway components mediate poised expression of the formative pluripotency programme in 2i.

Clustering based on the expression of ten naïve pluripotency marker genes showed that the downregulation of the naïve pluripotency TFs during formative differentiation is defective across multiple KOs (Figure 2a). Although expression of the naïve TFs was highly correlated, *Klf2* appeared to be an exception. *Klf2* downregulation was notably impaired in *Tcf7l1* KO ESCs, whereas it was unaffected by several KOs, including *Jarid2,* despite a comparable extent of deregulation of most other naïve marker genes. This indicates that *Klf2* expression can be uncoupled from the core naïve network. Forced *Klf2* expression stabilizes self-renewal ^24, 25^. *Klf2* depletion destabilizes mouse ESC identity ^26^ and increases the speed of Rex1-GFP downregulation upon 2i withdrawal (Supplementary Figure 2c). The differentiation delay in *Tcf7l1* KO ES cells is partially dependent on *Klf2* (Figure 2b), consistent with direct regulation^27^. Conversely, *Klf2* depletion in *Jarid2* KOs did not restore differentiation timing, indicating that separable gene-networks contribute to dismantling naïve pluripotency.

### Effects of exit gene depletion on global gene regulation

We used two measures to gauge differentiation delays in KOs compared to WT cells: first, the average expression levels of a set of naïve marker genes (defined throughout this manuscript as the average deregulation of the naïve marker genes *Esrrb, Nanog, Tfcp2l1, Tbx3, Prdm14* and *Klf4* at N24); second, the extent of global transcriptome adjustments between 2i and N24. Generally, the delayed extinction of the naïve TF network in KOs at N24 was accompanied by reduced gene expression change between 2i and N24 (R = −0.91; Figure 2c). However, some KOs deviated from this pattern. *Alg13, Jmjd1c* and *Jarid2* mutants showed a less pronounced change in naïve marker gene down-regulation than expected based on the reduced level of global gene expression adjustments towards an N24 profile. *Trim71* KO cells in contrast displayed modest global transcription changes but a more evident effect on naïve TF expression levels. Thus, the RNA binding protein Trim71 appears to be focused on the regulation of naïve pluripotency TF genes, although the ensuing exit delay is modest.

A general correlation was observed between the numbers of genes deregulated in 2i and at N24 (Figure 2d). Mutants with the strongest impact on the transcriptome (e.g. *Csnk1a1, Eed, Suz12, Zfp281 and Smg5*) showed deregulation of several thousand genes both in 2i and at N24. *Fgfr1* and *Tcf7l1* KOs were two exceptions to this correlation with relatively few genes deregulated in 2i, but several hundred at N24. We surmise that the effect of these two genes in 2i is largely masked because they are in the same pathways as the inhibitor targets Mek1/2 (PD0325901) and Gsk3 (Chiron 990201) ^7, 23^. Defective differentiation could simply equate with the extent of overall gene deregulation. We therefore mapped the average deregulation of naïve marker genes onto Figure 2d. Most KOs showing the strongest deregulation of naïve marker genes correlated with large-scale gene deregulation (both in 2i and at N24). However, there were several exceptions: for example, KO of *Eed*, *Suz12*, *Trim71*, and *Ctbp2* affected a substantial number of genes, but caused naïve marker deregulation that was weaker than expected assuming a direct correlation between the number of deregulated genes and the differentiation phenotype. Vice versa, KO of *Mbd3, Cdk8* or *Pten* showed marked naïve marker downregulation delays, despite a relatively mild impact on overall transcription (Figures 2d and Supplementary Figure 2d).

### Transcriptome analysis reveals a genetic interaction between *Jmjd1c* and *Tcf7l1*

The putative histone 3 lysine 9 (H3K9) demethylase *Jmjd1c (Kdm3c)* KO is a case of particular note. Few genes were significantly deregulated at N24 (Figure 2d), with a mild global neutralisation of differentiation-induced transcriptome changes (Figure 2c). Neither the role of *Jmjd1c* in pluripotency regulation, nor its mode of action are known. We used multiple regression analysis to determine the similarity of the *Jmjd1c* KO to all other KO RNA-seq profiles. At N24, the *Jmjd1c* KO transcriptome was most similar to the *Tcf7l1* KO profiles (Supplementary Figure 2e), suggesting a potential functional connection between *Jmjd1c* and *Tcf7l1*. Indeed, Jmjd1c has recently been reported as a high-confidence protein-interactor of Tcf7l1 ^28^. The 2i component Chiron 99021 is a specific inhibitor of Gsk3 and phenocopies deletion of *Tcf7l1*, which is the downstream repressor in ESCs ^29, 30^. Accordingly, Chiron 99021 delays the differentiation of WT ESCs, but had little additional effect on *Tcf7l1* KOs (Figure 2e). *Jmjd1c* deficient ESCs did not show a discernible phenotype at N72, in line with minimal deregulation of the naïve TF network. Addition of Chiron 99021 had a stronger than expected effect on Jmjd1c KOs and resulted in a synthetic enhanced delay phenotype (Figure 2e), suggesting a cooperative activity of *Jmjd1c* and the Wnt/Tcf7l1 axis in the exit from naïve pluripotency.

### Relative quantification of differentiation delays *in vitro*

To quantify differentiation delays, we obtained RNA-seq data from a 2h-resolved WT ESC differentiation time course (Figures 3a and Supplementary Figure 3a). We then compared the expression patterns of specific gene-sets in a given KO at N24 to the expression of the same gene-set along the WT 2i to N32 trajectory. This enabled us to ‘position’ each KO along the trajectory and thus quantify the differentiation delay with a precision of about 4h (Figure 3b, Supplementary Table 2). In a complementary approach, we used all 3058 DEG between 2i and N24 as an alternative reference set. This yielded similar, but on average slightly less pronounced differentiation delays (Figures 3c and Supplementary Figure 3b), supporting the hypothesis that the naïve TF network is regulated in concordance with but partially independently from the rest of the transcriptome during exit from naïve pluripotency.

**Figure 3.**
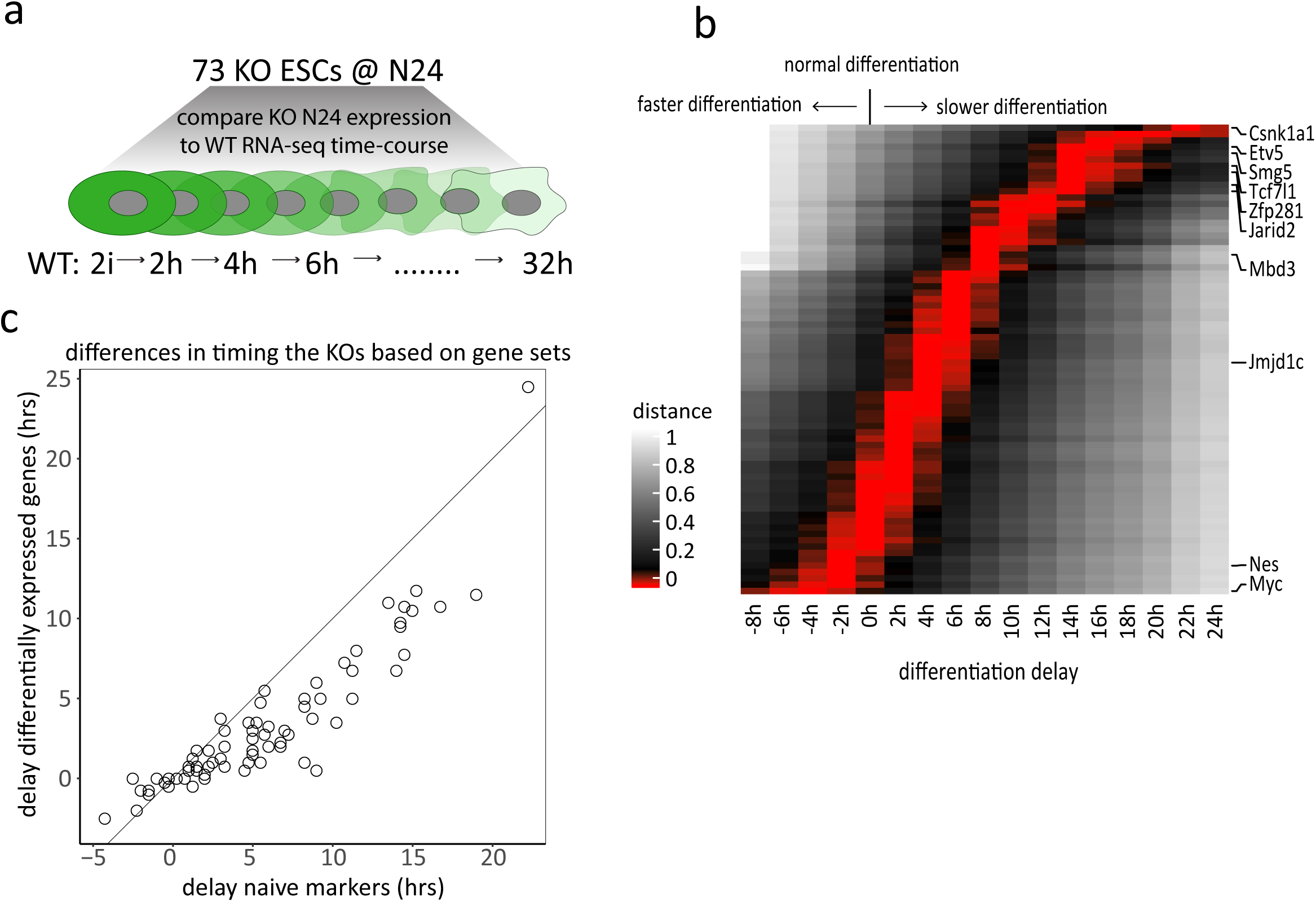
**a,** Scheme illustrating comparisons of the 73 KO gene expression profiles at N24 with a 2h-resolution *in vitro* differentiation time course of WT RC9 ESCs. **b,** Heatmap showing differentiation delays of the 73 KO lines quantified by average naïve marker gene expression. Red bars indicate closest correlation to that specific time-point. Positive values indicate delayed differentiation. Negative values more rapid differentiation. Each line corresponds to one KO. Selected KOs are indicated. See Supplementary Table 2 for the full hierarchy of the 73 KOs. **c,** Plot comparing the differentiation delays calculated using naïve markers as in **b,** or using all 3058 genes changed in WT differentiation from 2i to N24.

### The regulatory program for pre- to post-implantation epiblast transition is preserved in most mutants

To explore the extent to which our *in vitro* data captures *in vivo* regulation, we first compared global transcriptomes of WT and KO ESCs to single cell transcriptome data from the *in vivo* pre- (E4.5) and post-implantation (E5.5 and E6.5) mouse epiblast ^10^ (Figure 4a,B). As expected ^8^, WT and KO cells in 2i showed greater similarity (fraction of identity - FOI) to the E4.5 than to the E5.5 epiblast. Cells at N24, in contrast, showed higher FOI to the E5.5 epiblast. Cells in 2i and at N24 showed low FOI to E6.5 epiblast, underscoring that 24h 2i withdrawal models the E4.5 to E5.5 epiblast transition ^2^. Interestingly, some KO ESCs showed expression features in 2i with increased similarity to the *in vivo* epiblast (Figures 4c,d and Supplementary Table 3). Among those were KOs showing strong *in vitro* differentiation defects, such as *Zfp281, Tsc2* and *Etv5,* but also *Trim71* and *Smg7*, which showed only a modest differentiation delay in culture.

**Figure 4.**
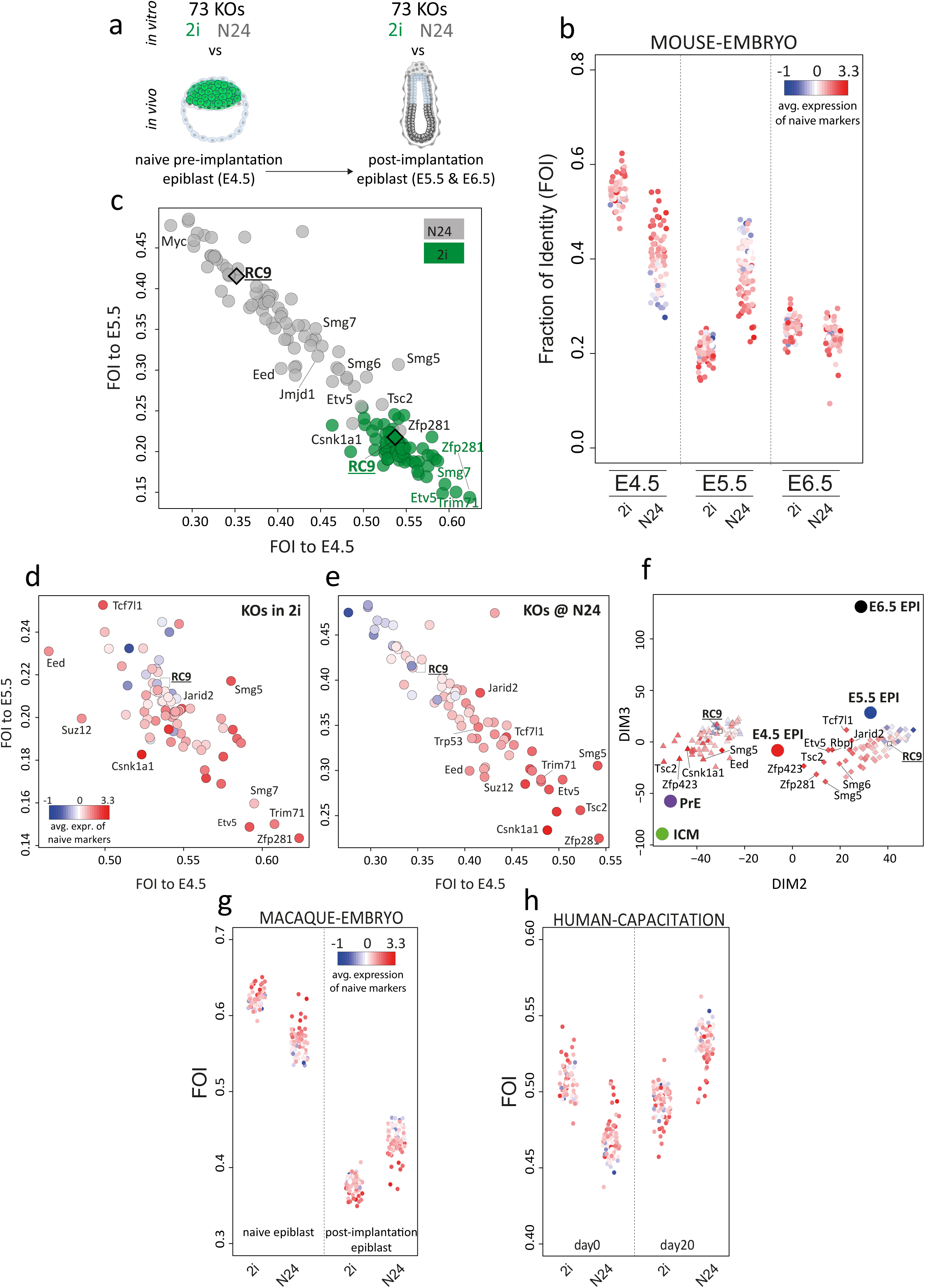
**a,** Scheme illustrating the comparison of gene expression profiles of 73 KOs in 2i and at N24 with *in vivo* derived averaged single cell expression data comprising peri-implantation epiblast tissues. Specifically, transcription profiles of each KO in 2i and at N24 were compared with in expression datasets obtained from the E4.5, E5.5 and E6.5 epiblast. **b,** Fraction of identities (FOI) between 2i and N24 expression data and mouse E4.5, E5.5 and E6.5 epiblast, computed using all expressed genes (log2FPKM > 0) for all KOs. Each dot represents one KO. Average naïve marker expression in KOs at N24 is indicated by a colour gradient. **c,** Plot showing a comparison of FOIs of WT (RC9) and KO cells in 2i (green) and at N24 (grey) to E4.5 and E5.5 mouse epiblast, computed using all expressed genes (log2FPKM > 0). **(d,e)** Data shown in Figure 4c separated by culture condition. Comparison of FOIs of WT and KO cells in 2i **d,** and at N24 **e,** to E4.5 and E5.5 mouse epiblast; average naïve marker expression in KOs at N24 is indicated by a colour gradient. **f,** PCA showing all KOs in 2i (triangles) and at N24 (diamonds) and embryo derived datasets (ICM, PrE, E4.5Epi, E5.5Epi, E6.5Epi). As in **d,**, blue to red colour gradient indicates naïve marker expression at N24 of the respective KO. **g,** Similar to Figure 4b, showing FOIs between 2i and N24 datasets and macaque pre- and post-implantation embryos. **h,** Similar to Figure 4b, showing FOIs between 2i and N24 datasets and hESC capacitation datasets comprising hESCs in naïve conditions (day0) and hESCs 20 days after release into primed conditions (day20).

At N24 several KOs, such as *Zfp281*, *Tsc2*, *Smg5* and *Smg6*, retained strong similarity to the E4.5 epiblast (Figures 4c,e and Supplementary Figure 4a). Strikingly, *Zfp281* KO profiles at N24 showed similarity to the E4.5 epiblast on par with WT cells cultured in 2i (Figures 4c and Supplementary Figure 4a), consistent with an overt differentiation delay phenotype ^31^.

Overall, there was a good correlation between the *in vitro* differentiation delay and the similarity with pre-implantation epiblast cells: the N24 KO transcriptomes that were more similar to the E4.5 epiblast exhibited stronger exit-delay phenotypes (Figure 4e, Supplementary Figure 4b). Together, this indicates that similar transcriptional networks are regulated by similar mechanisms during *in vivo* and *in vitro* transitions to formative pluripotency. However, at N24 three KOs (*Jarid2*, *Tcf7l1*, and *Trp53*) showed a similarity to the E4.5 epiblast which was smaller than expected based on their *strong in vitro* differentiation defects (Figures 2a and 4e).

We then performed principal component analysis (PCA) for genes variably expressed in embryo development, using averaged values for inner cell mass (ICM), primitive endoderm (PrE) and E4.5, 5.5 and 6.5 epiblast ^10^. The PCA separated the 2i and N24 transcriptomes into two clusters with proximity to PrE/EPI and E5.5 respectively. Notably*, in vitro* differentiation defects were reflected by closer proximity of mutant N24 profiles to the E4.5 epiblast (Figure 4f). Overlaying the PCA with a colour gradient indicating similarity to PrE shows that multiple KOs in 2i gained similarity to PrE (Supplementary Figure 4d). Concordantly, the expression of PrE-defining transcription factors *Gata4*, *Gata6*, *Sox7* and *Sox17,* was readily detectable in *Eed, Suz12, Zfp423* and *Csnk1a1* KOs in 2i (Supplementary Figure 4e). A shift toward PrE-identity was most pronounced for the PRC2 KOs *Eed* and *Suz12*. Notably, this group of KOs (Csnk1a1, PRC2, Zfp423, which exhibited substantial *in vitro* differentiation defects while showing reduced similarity to the E4.5 epiblast, also showed decreased similarity between their N24 profiles and the E5.5 epiblast (Supplementary Figure 4d). We therefore suggest that deregulation of the PrE-programme above a certain threshold has disengaged these KOs from the normal embryonic developmental trajectory. In contrast KOs showing relatively increased similarity to the E4.5 in 2i and decreased similarity to the E5.5 epiblast at N24 are deficient for genes regulating the formative transition during development (e.g. *Zfp281*, *Etv5*, *Rbpj* and, *Trim71*, Supplementary Figure 4d).

### Correspondence to primate embryogenesis

Despite morphological and timing differences between rodent and primate peri-implantation development, embryos of both orders appear to transit through similar pluripotency states ^1, 32^. To examine this issue, we compared transcriptional profiles of our KO series and cells of the macaque *in vivo* naive- and post-implantation epiblast ^33^. In general, ESCs in 2i were more similar than cells at N24 to the pre-implantation macaque epiblast (Figure 4g). Correspondingly, N24 cells were closer to the macaque post implantation epiblast. Interestingly, KOs showing higher identity with the mouse *in vivo* epiblast also displayed an increased similarity with the macaque pre-implantation epiblast (Supplementary Figures 4f,h). We also compared our datasets to data obtained from *in vitro* capacitation during human naïve to primed ESC differentiation ^34^. KOs displaying strong differentiation defects in mouse cell culture had higher correspondence to naïve human ESCs and, at N24, lower correspondence to primed human ESCs (Figures 4h and Supplementary Figure 4g,I). These observations suggest that the overall GRN-redeployment during the naïve to formative transition is conserved between rodents and primates.

### Identification of an extended naïve pluripotency network

ESC differentiation requires fine-tuned coordination between extinction of the naïve and initiation of the formative transcription networks. To date, only incomplete inventories of the genes that functionally define these two states have been made. These genes cannot be defined based simply on differential expression between 2i and N24, because not all DEG will be functionally linked to the naïve GRN. To identify those genes that show specific linkage to the core naïve pluripotency network, we trained regression models to predict expression changes across all KOs (in 2i and at N24) as a function of naïve marker log-fold-changes. The 374 genes whose expression was tightly correlated (R^2^ > 0.7) with one or several of the seven core pluripotency markers (*Nanog, Esrrb, Tbx3, Tfcp2l1, Klf4, Prdm14 and Zfp42)* were defined as ‘Naive-Associated Genes’ (NAGs; Supplementary Table 3). The naïve pluripotency TFs not represented in the defining TF-set were correctly identified as NAGs, including *Klf5, Nr0b1* and *Nr5a2*. However, *Klf2* is not one of the NAGs being only weakly associated (R^2^=0.53) with the naïve network. This concords with our earlier observation that *Klf2* expression can be uncoupled from the naïve TF network. Of further note, *Klf2* expression is barely detectable in marmoset or human pre-implantation epiblast cells (Supplementary Figure 5b).

The NAG showing the highest correlation with the naïve core network is *Pdgfa*, which is relatively highly expressed in ES cells (FPKM ∼70). *Pdgfa* has no known role in ESC self-renewal, but functions in segregation of the primitive endoderm ^35^. To examine whether the link between *Pdgfa* and the naïve transcription factor network is maintained *in vivo*, we utilized GRAPPA, a tool to visualize single cell expression data from preimplantation development ^36^. Indeed, *Pdgfa* is uniformly expressed in the E4.5 epiblast (Supplementary Figure 5c). Interestingly, its cognate receptor *Pdgfra* is neither expressed in ES cells nor the naïve epiblast (FPKM in WT ESCs <0.5), but specifically marks the neighbouring primitive endoderm at E4.5 ^37^.

We surveyed the expression of NAGs at the transition from naïve to post-implantation pluripotency in the single cell RNA-seq datasets from E4.5, E5.5 and E6.5 epiblast cells ^10^. We detected a clear enrichment of NAGs within genes that separate the pre- from the post-implantation epiblast in the differentiation-state resolving dimension of a principal component analysis (PCA) (Figures 5a and Supplementary Figure 5d), highlighting that NAGs are a set of indicators for the naïve epiblast state *in vitro* and *in vivo*. Strikingly, NAGs showed a near identical fold change behaviour *in vitro* and *in vivo* during the epiblast transition from E4.5 to E5.5 (Figure 5b,e,f). Notably, this effect was not observed for the top 250 differentially expressed genes in ESC differentiation *in vitro*. Furthermore, NAG orthologues showed significantly co-regulated expression dynamics during macaque pre- to post-implantation epiblast differentiation *in vivo* (Figure 5c,e,f) and during capacitation of human ESCs *in vitro* (Figures 5d,e,f), which further underscores their relevance. We thus propose that NAGs constitute an integral component of the naïve transcriptional network with potential functional relevance *in vivo* across mammalian species.

**Figure 5.**
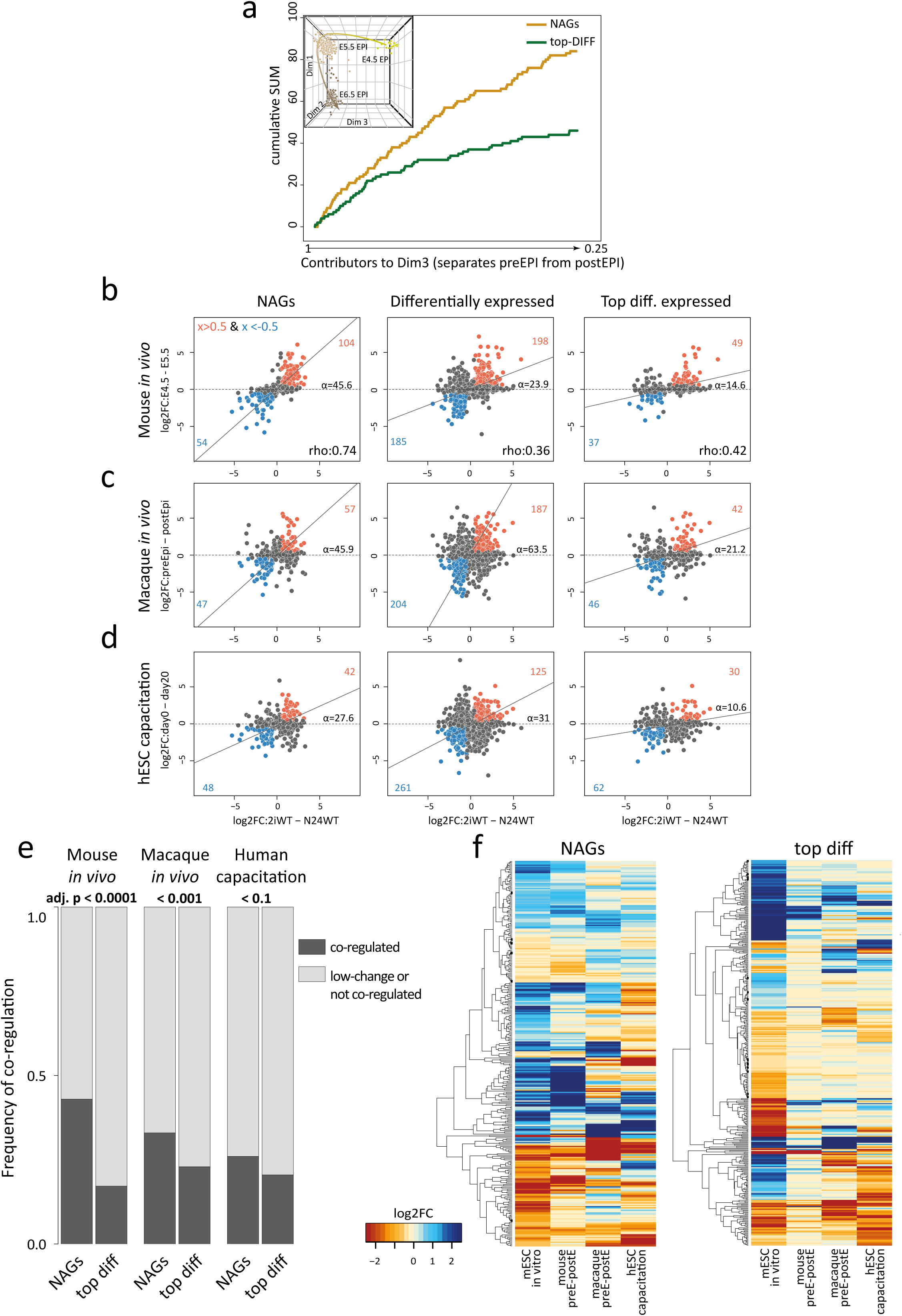
**a,** x-axis displays all 3^rd^ dimension (Dim3) genes, which separates mouse naïve from post-implantation epiblast (the inset shows the full PCA plot, see also Supplementary Figure 5d) in order of extent of contribution to Dim3. The contribution to Dim3 of NAGs (orange) and the top 250 differentially regulated genes (green) are plotted in a cumulative manner. **b,** Plots comparing relative log2FC in mouse ESC differentiation (2i vs N24) to relative log2FC in mouse *in vivo* transition from pre- to post-implantation epiblast (E4.5 vs E5.5). Selected gene groups are plotted: i) NAGs, ii) all 3058 differentially expressed genes in mESC differentiation and iii) the top 250 up and downregulated genes in mESC differentiation. Rho values indicate the level of correlation between *in vivo* and *in vitro* differentiation of given gene groups. Alpha indicates the angle between the x-axes and the orthogonal regression. Genes with log2FC > 0.5 and < −0.5 in both the x- and y-axes are highlighted in orange and blue respectively. **c,** As for **b,** except plots comparing log2FCs in mouse ESC differentiation (2i vs N24) to macaque *in vivo* transition from pre- to post-implantation epiblast cells. **d,** As for **b,** except plots compare log2FCs in mouse ESC differentiation (2i vs N24) to human naïve ESC capacitation (d0 to d20). **e,** Graphs showing the frequency of co-regulation of NAGs or the top 250 differentially expressed genes (2i vs N24) with differentially expressed genes in mouse and macaque *in vivo* pre- to post-implantation epiblast differentiation, and human naïve ESC capacitation. Two-tailed chi-square test (CI 95%) was used to compute significance levels between the expected and the observed number of modulated genes. **f,** One-way hierarchical clustering showing expression changes of NAGs (left) and top differentially expressed genes (right) in the indicated differentiation models.

### Deregulation of signaling cascades is a hallmark of differentiation delay

We next analysed the extent of deregulation of five key signalling pathways known to be active in pluripotent cells: LIF/Stat3, mTOR, Wnt/β-catenin, Fgf/ERK and Notch ^7, 38–42^. Changes in the activity of a signalling pathway are not necessarily reflected in expression changes of pathway member transcripts. Thus, in order to quantify pathway activities, we employed ‘expression footprints’. We identified pathway specific marker gene-sets, each containing 50 genes, reporting pathway activity changes. These marker sets were determined using N24 derived KO transcription-profiles of ESC lines deficient for key signalling pathway components. Thereby we defined key mTOR (affected by *Tsc2 KO*), Wnt/β-catenin signaling (*Tcf7l1 KO*), Fgf/ERK (*Fgfr1* and *Ptpn11 KOs*), and Notch (*Rbpj KO*) pathway targets (Supplementary Figure 6a, Supplementary Table 4). Overlaps with available Tcf7l1 and Rbpj ChIP and activity profiles ^27, 43^ supported the reliability of this approach (Supplementary Figure 7E). For LIF-signaling, we compared RC9 ESCs grown in 2i in the presence and absence of LIF for 24 hours. This resulted in a list of LIF-sensitive genes including known targets like *Socs3, Gbx2, Junb, Tfcp2l1, Klf4* and *Klf5* ^27, 44^ (Supplementary Table 4). In summary, the expression footprints present non-overlapping sets of marker genes whose expression state is indicative of the activity of the respective signaling pathway.

Using the expression footprints, we then asked whether we could detect a preferential deregulation of one or more of these signalling footprints in specific KOs either in 2i (for LIF) or at the N24 time-point (for all other pathway-profiles) (Figure 6a, Supplementary Table 4). Surprisingly, we detected mis-regulated LIF target genes in several KO ESC lines cultured in 2i in the absence of LIF (Figure 6a). Activation of such a ‘LIF-like‘-profile in 2i was closely correlated with the extent of differentiation delay observed at N24 (Supplementary Figure 6b). Notably, presence of LIF together with 2i before induction of differentiation slows down naïve state exit (Supplementary Figure 6c) ^4, 16^. *Ptpn11*, *Zfp281*, *Tsc2* and *Trim71* KO ESCs showed the greatest similarity to the LIF profile in 2i; in contrast, *Tcf7l1* KO ESCs, despite showing a pronounced differentiation defect, lack the ‘LIF footprint’ (Supplementary Figure 6b). Addition of a JAK inhibitor to several KOs showing a LIF-like profile resulted in no or only minor amelioration of the differentiation defect. This suggests that these KOs do not directly activate Jak/Stat signalling, but that a LIF-like expression profile reflects a consolidated naïve network that is resilient to dismantling.

**Figure 6.**
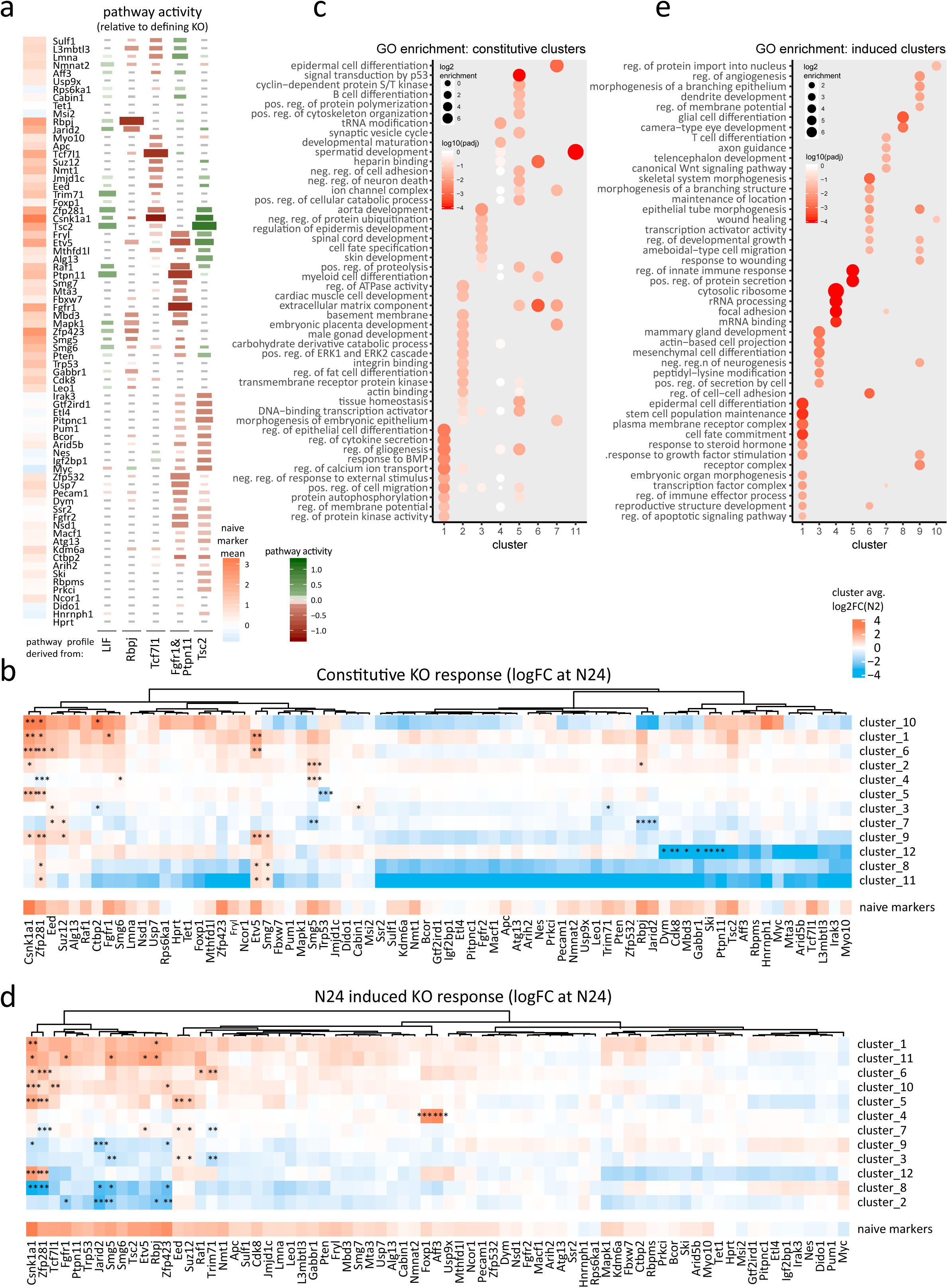
**a,** Relationship between the KO expression profiles at N24 (or in 2i for the LIF pathway) to the expression footprints of the five signalling pathways as indicated (see also Supplementary Figure 6a). Relative pathway activity is −1 for Fgf/ERK (defined by Fgfr1 & Ptpn11 KOs), Wnt (Tcf7l1 KO) and Notch (Rbpj KO) and +1 for mTOR (Tsc2) and LIF (defined by presence of LIF in 2i medium) for effects equally strong as in the representative KO. Pathway activity is indicated by colour code (green for activity, red for inactivity). All tiles with an absolute pathway activity smaller 0.3 are grey. The size of the tiles corresponds to the spearman correlation of pathway footprints between each KO and the corresponding representative KOs. The colour gradient on the sidebar at the left indicates mean change of naïve marker gene expression (as a measure of phenotypic strength) for the corresponding KO at N24. **b,** Heatmap showing average log2FCs (KO^N24^ vs. RC9^N24^) of constitutive response gene clusters. Asterisks indicate that the absolute row-wise z-value of the respective mean expression value was above 1.96 (p ≤ 0.05), indicating extreme values within the context of that row. For reference, average naive marker log2FCs at N24 vs WT are shown in a separate row below (similar to Figure 2a). **c,** GO enrichment analysis per ‘constitutive KO response’ cluster. Dot-size scales with log2 enrichment. Adj. p vales are clour coded in shades of red. **d,** As for **b,** except heatmap showing average log2FCs (KO^N24^ vs. RC9^N24^) of N24 induced KO response gene-clusters. **e,** As for **c,** except GO enrichment analysis per ‘N24 induced KO response’ cluster.

As expected, *Raf1, Fgfr2, Mapk1* and *Etv5* KO ESCs showed an Fgf-signalling footprint at N24. *Mbd3*, *Mta3*, *Nsd1* and *Arid5b* KOs showed Fgf/ERK target deregulation similar to reference KOs, indicating an involvement of chromatin regulators in ERK target-gene control (Figure 6a). *Tsc2* deficiency lead to constitutive activity of the mTORC1 pathway, which has previously been associated with an exit from naïve pluripotency phenotype ^45^. Many KOs (including *Pten*) exhibited deregulated Tsc2 responsive genes, suggesting that regulation of mTORC1 (or its downstream effectors) is a common node to gate the exit from naïve pluripotency. *Tsc2* depletion resulted in partial activation of a LIF footprint, suggesting that a component of the LIF response is mediated through Akt ^39, 46^. *Rbpj* and *Tcf7l1* KOs did not show a correlation with each other or with Tsc*2* KO, consistent with independent mechanisms ^21^. The beta-catenin destruction complex member *Apc* and the previously discussed *Jmjd1c* showed the expected similarities with the *Tcf7l1* profile. Interestingly, several KOs with strong phenotypes showed footprints similar to *Tcf7l1* KOs at N24. This is in line with evidence that β-catenin constitutes or regulates a major differentiation switch during the exit from naïve pluripotency and is influenced by multiple exit KO genes ^29, 47^. Genes downstream of the Notch pathway were most strongly affected in a group of mutants deficient for mRNA homeostasis (*Smg5* and *Smg6*) and chromatin regulators, including *Jarid2, L3mbtl3 and Mbd3*. The latter suggests an interaction between the Notch pathway (*Rbpj*) with the Polycomb and Nurd complexes to modulate network rewiring during the exit from naïve pluripotency. Cooperativity between Rbpj and the Polycomb-associated protein L3mbtl3 has been reported in *Drosophila* and *C. elegans* ^48^.

We asked to what extent the alteration of any of the five signalling pathways was predictive for the strength of the differentiation defect. We found that whereas aberrant Fgf/ERK or mTORC1 activity were unreliable predictors, transcriptional similarity to *Rbpj* or *Tcf7l1* KOs at N24 or deregulation of LIF target genes in 2i correlated with differentiation delay (Supplementary Figure 6d). These data indicate that many KOs with a delay phenotype have altered activity of at least one of the five key signalling pathways. Strikingly, however, there was no single signalling pathway perturbed downstream of all KOs. We observed that KOs showing multiple pathway footprints are rare, but most KOs showing a delayed naïve marker gene downregulation at N24 deregulate at least one specific pathway. *Trp53*, *Ncor1* and *Usp9x* constitute exceptions, which show appreciable differentiation defects without deregulating of any of the tested signalling cascades. We conclude that signalling pathways are harnessed in parallel and that most KO genes contribute directly or indirectly to at least one pathway activity that promotes the exit from naïve pluripotency.

### Identification of gene-networks downstream of multiple KO genes

To explore common regulatory targets of multiple KO-factors, we classified deregulated genes into two groups: (i), deregulated by a KO in both 2i and at N24, termed ‘constitutive KO response’; (ii) not, or only weakly, deregulated by a KO in 2i, but significantly deregulated at N24, termed ‘N24 induced KO response’. Genes in both groups are affected by knockouts, but genes in the second group deviate from wild-type expression only under conditions of differentiation. We identified all genes that showed either type of behaviour in at least one KO (Supplementary Figure 6e; Supplementary Table 5). Genes deregulated exclusively in PRC2 core-components (*Eed* or *Suz12* KOs) or *Csnk1a1* KOs were excluded from further analysis, because the deregulation of thousands of genes in these KOs had a disproportionate impact on the resulting gene-lists. In total, the constitutive and the N24 induced KO response groups contained 1,837 and 720 genes, respectively. Genes belonging to the N24 induced KO response group were strongly enriched for factors that are dynamically regulated during normal differentiation, whereas genes of the constitutive response group were not (Supplementary Figure 6f). Thus, naïve marker genes were absent from the constitutive group. Together with NAGs, they were strongly enriched in the N24 induced group, because deregulation of the naïve network only initiates upon 2i withdrawal.

To characterize the constitutive KO response genes, we grouped them based on their log-fold-changes between KO and WT at N24. This resulted in twelve clusters, ranging in size from 20 to 597 genes (Supplementary Table 5) (Figure 6b). By Gene Ontology (GO) analysis, eight of the twelve clusters showed a significant enrichment of at least one GO term containing more than five genes (Figure 6c, Supplementary Table 5). Many of these GO terms relate to cell fate specification and development. Notably, the constitutive response clusters often shared identical or similar GO terms, suggesting functional convergence. NAGs were underrepresented in constitutive clusters and only contained in clusters 1 (const c1) and 2, for which expression profiles across all KOs correlated with naïve marker expression (Figure 6b). Therefore, deregulation of genes within const c1 and c2 in 2i is proposed to directly translate to differentiation defects observed at N24. Cluster 1 genes were enriched for general differentiation-related factors and response to Bmp signalling. Constitutive cluster 2 contained a significant number of Fgf/ERK pathway defining genes (Supplementary Figure 7e) and was consistently enriched for Fgf/ERK signalling related terms (Supplementary Table 5). Intriguingly, constitutive cluster 10 contained multiple *Zscan4* isoforms ^49^ and 14 out of 45 genes in this cluster are bound by the co-repressor Trim28, which is involved in silencing of endogenous retroviruses ^50–52^. Strong upregulation of this cluster was observed in *Csnk1a1* and *Zfp281* KOs, but strong downregulation in *Rbpj* and *Jarid2* KOs. Overall, cluster 10 regulation showed no strict correlation with a differentiation delay phenotype.

Analysis of the N24 induced response genes yielded 12 clusters ranging from 8 to 103 genes (Figure 6d and Supplementary Table 5). All naïve marker genes were detected in N24 induced cluster 1 (N24i c1), with the notable exception of *Klf2*. *Klf2* was part of N24 induced cluster 10, within which more than half the genes are Tcf7l1 ChIP targets ^27^ (Supplementary Figure 7E). This provides further support for the notion that Wnt/Tcf7l1 controls *Klf2* expression independently of the rest of the naïve network. Interestingly, twelve percent of genes in N24 induced cluster 10 were screen hit genes, suggesting a major impact of this cluster on the exit from naïve pluripotency (Supplementary Figure 7e). NAGs were distributed in eight out of the twelve N24 induced clusters. In marked contrast to the constitutive KO response, individual N24-induced clusters displayed specific functional enrichments (Figure 6e). Therefore, the N24 induced response clusters represent distinct molecular pathways and cellular functions that were altered in multiple KOs. N24 induced cluster 7 was significantly regulated in *Etv5*, *Trim71* and *Zfp281* KOs at N24 (Figure 6d). Cluster 7 contains pivotal cell fate switch genes like *Fgf4*, *Tfe3* ^14^ and *Tcf7l1* itself, but NAGS were underrepresented in this cluster. Notably, *Etv5, Trim71* and *Zfp281* are the three KO lines that show the strongest exit delay while retaining E4.5 identity at N24 (Supplementary Figure 4d), suggesting that N24 induced cluster 7 contains crucial cell fate switch genes that are modulated during *in vitro* and *in vivo* exit from naïve pluripotency. In summary, constitutive and N24 induced clusters define crucial genetic modules that are co-regulated by multiple exit factors.

We asked whether the expression status of these genetic modules correlated with the activity of differentiation regulating pathways. We utilized pathway activity levels and cluster specific expression levels across all KOs to build a model predicting the impact of pathway activity on cluster-gene expression (Figures 7A and S7A to F). Adding further evidence to the pathway to cluster relationships, we found that in most cases activity changes of the regulatory pathways preceded the expression changes of their putative target clusters over the 2h differentiation time course. The Notch and mTor pathways showed high connectivity to N24-induced clusters (8 and 6 out of 12 clusters, respectively), whereas Wnt/βcatenin signalling was specifically correlated with clusters containing high levels of NAGs (N24i c1 and c10) and the only constitutive cluster containing an appreciable number of NAGs (const c1) (Figure 7A). Together, this shows that at least 24 distinct transcription modules can be defined, whose expression is deregulated upon exit gene depletion and which are largely under the control of five key signalling pathways gating differentiation.

**Figure 7.**
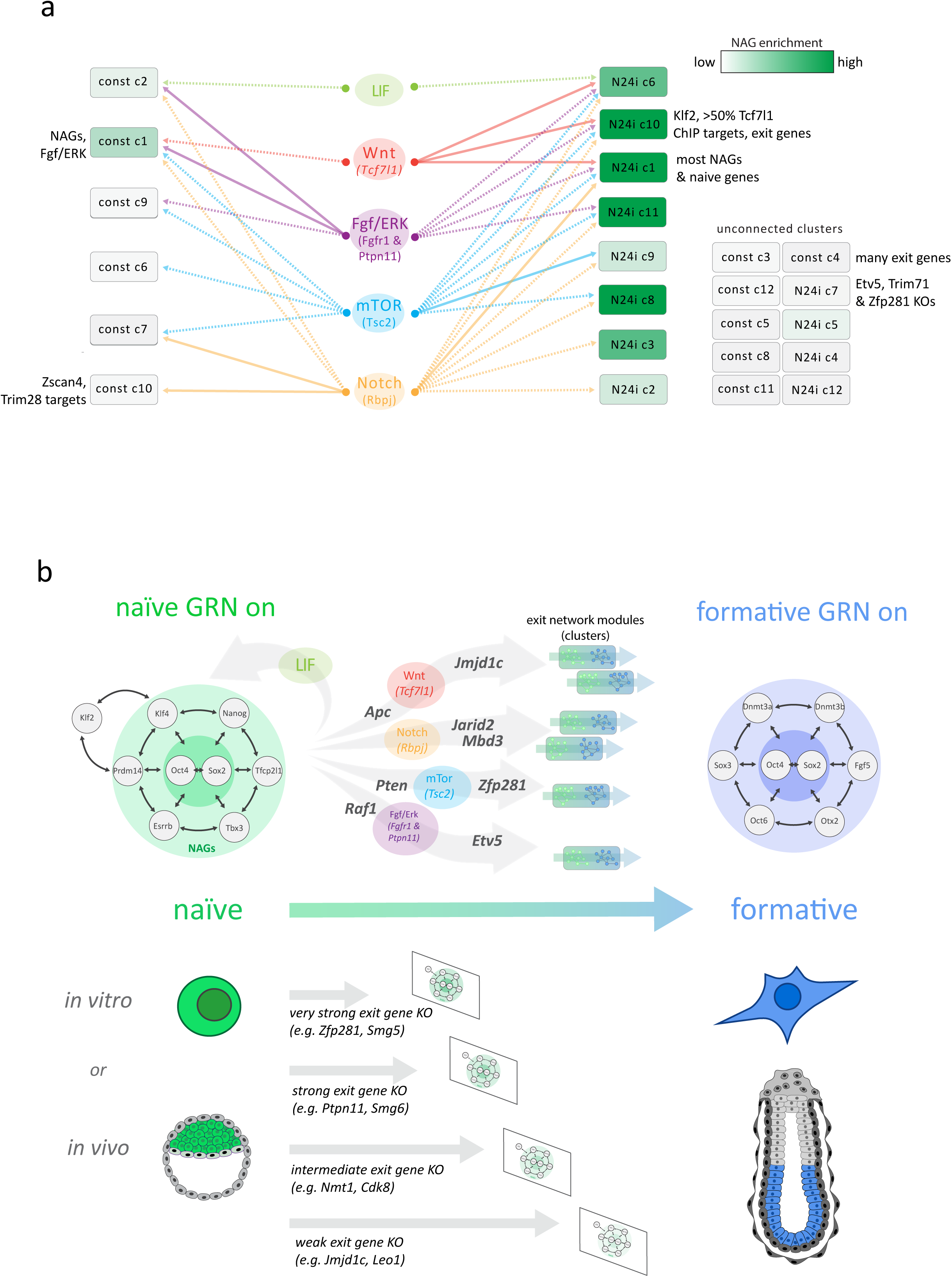
**a,** Visualization of connections between pathways (central ellipses) and gene clusters (rectangles) based on multiple regression models. The colour gradient of the clusters indicates enrichment for NAGs. Dashed connections link pathways to clusters that fit time-course validated regression models (see Supplementary Figure 7A and C). Continuous connections are further supported by ChIP or pathway defining gene enrichments (see Supplementary Figure 7E,F). Connections are coloured by source pathway. Unconnected clusters are listed at the right. **b,** Summary model of the genetic and signaling pathway control of naïve to formative transition. Between the naïve and formative gene regulatory networks, the five signaling pathways largely operate through partially overlapping separable gene clusters.

## Discussion

Ordered progression through pluripotency requires shutdown of the naïve TF-network and concomitant large-scale rewiring to establish the formative GRN ^2, 3, 19^. Following Harald Weintraub’s pioneering work with muscle differentiation and Thomas Graf’s work on haematopoiesis, the ability of certain transcription factors to change cell identity without a developmental context gained central prominence with somatic cell reprogramming to pluripotency ^53–55^. However, most of these studies involved forced expression of selected transcription factors to achieve synthetic cell state transitions *in vitro*. Here we utilized the remarkable properties of ESCs to recapitulate the pivotal embryonic transition from naïve to formative pluripotency. Notably, we show that this developmental cell state transition is not based on the action of one or two master transcription factors. Although crucial for defining lineage trajectories, transcription factors do not initiate cell fate decisions. Instead, the robust stability of naïve pluripotent stem cells is sustained by multiple crosstalking components and is dismantled by at least four diverse signalling inputs.

Our dissection of an authentic cell state transition indicates that the exit from naïve pluripotency is largely guided along a trajectory constrained by signalling pathways and funnelled through a handful of genetic modules. These adaptations of cellular networks establish the opportunity for TFs to initiate and consolidate cell identity. In the specific case of the exit from naïve pluripotency, expression of Oct4 and Sox2, which are key TFs for both the naïve and formative states, provide cornerstones for both the naïve and formative regulatory networks to ensure lineage fidelity during differentiation (Figure 7B). Our data support a cell fate transition model in which extinction of cell identity induced by the interplay of signalling and the chromatin modification machinery precedes the initiation of a novel cell fate.

Several signalling-cascades have been implicated in ESC self-renewal and naïve pluripotency exit, including pathways controlled by LIF, Akt/mTOR, Wnt, Fgf/ERK and Notch. We derived transcription footprints of these pathways and scrutinized the extent to which they were deregulated by the different KOs to find that virtually all KOs showing a strong differentiation defect also showed deregulation of at least one of these five pathways. Notably, however, we observed segregation in terms of pathway deregulation. Thus, KOs that induce e.g. a *Tcf7l1*- like profile were less likely to show a signature of another pathway. This suggests that ESC differentiation requires coordinated activity changes of independent pathways under control of separable genetic networks. The observation that single gene depletion is not sufficient to prevent exit from the naïve state is consistent with this interpretation ^21^. Our analyses exposed discrete pathway and GRN features that mediate timely and robust mammalian cell state transitions. We found that 12 genetic sub-networks (clusters) with non-overlapping cellular functions emerge during the exit from naïve pluripotency. Interestingly, signalling pathway activity was closely co-regulated with these clusters suggesting a functional linkage. This implies that formative differentiation is driven by a limited number of parallel acting genetic modules under control of, or controlling, key signalling pathway activities.

By examining co-regulation with the core naïve network across 146 perturbations (73 KOs in 2i and at N24) we identified the NAGs. NAGs obey strikingly similar expression dynamics *in vitro* and *in vivo*. Moreover, in the macaque pre- to post-implantation transition and in human naïve cell capacitation, orthologous NAGs show similar behaviour as during mESC exit from naïve pluripotency. Therefore, this cohort of genes constitutes a layer of the pluripotency network that is tightly linked to, and likely acts in conjunction with, the core naïve TF-network in mammals ^4^. We suggest that collective modulation of NAG expression will propel naïve cells into formative differentiation and conversely, that collective NAG deregulation will delay proper differentiation. The control of NAG expression appears to be interwoven with the signalling pathways involved in controlling the naïve to formative transition. The N24 induced clusters enriched for NAGs showed high connectivity to all tested pathways. The functional significance of NAGs likely extends beyond self-renewal and naïve identity. Indeed, most NAGs are not transcription factors. The growth factor Pdgfa is a case in point. *Pdgfa* is the most strongly associated NAG, but has no known activity on ESCs, which do not express the cognate receptor, Pdfgra. Within the ICM, Pdgfa expression is restricted to the naïve epiblast, whereas Pdgfra is present exclusively in the primitive endoderm. This reciprocal expression pattern is consistent with the known paracrine action of Pdgfa to promote primitive endoderm segregation ^35^. Thus, conserved linkage of NAGs to the naïve TF network may mediate paracrine communication to other lineages within the blastocyst as well as consolidate naïve epiblast fate.

Whereas most naïve TFs were tightly co-regulated across the 73 KOs, *Klf2* expression was uncoupled from other members of the core network. Interestingly, while *Klf2* is highly expressed in mouse ESCs and naïve epiblast, it is very lowly expressed in primate blastocysts, and also in the porcine pluripotent compartment ^56^. *Klf2* may therefore be a rodent specific addition to the naïve TF-network. A further significant species difference is the very low expression of *TCF7L1* in primate naïve cells, which underlies the differential responsiveness of mouse and human naïve cells to GSK3 or Wnt pathway inhibition ^34, 57^. *Klf2* is a genomic target of Tcf7l1 ^27^. Interestingly, *Klf2* and *Tcf7l1* are found in the same N24 induced gene-expression cluster (cluster 10), suggesting coregulation of both components of this mouse specific naïve pluripotency feedback loop. For most KOs the degree of failure in naïve TF downregulation and the neutralisation of global differentiation-related transcription changes are comparable. However, the relatively weak *Trim71* phenotype despite specific upregulation of some naïve TF-genes, suggests that up-regulation solely of core naïve factors must reach a certain threshold to stably maintain naïve pluripotency, consistent with negative feedback constraints within the network ^4, 5, 58^.

The remarkable advantages of ESCs as an experimental venue include the utilisation of well-defined culture conditions to recapitulate key features of *in utero* developmental progression ^1, 2, 59, 60^. Our high-resolution transcriptome analysis confirms that differentiation over 24 hours *in vitro* mimics the peri-implantation transition from naïve to formative epiblast *in utero.* These data provide rich resources for further studies. In particular, many strong differentiation delay KO transcriptome profiles retained transcriptome resemblance to the pre-implantation epiblast. This relationship held true in 2i and at N24 and indicates that the KO of these genes perturb the operative *in vivo* cell state transition machinery. In general, we surmise that stalling the activity of the exit machinery is critical to *in vitro* capture and self-renewal of naïve stem cells, consistent with empirical requirements for signal inhibition in both mouse and human ^7, 57^. Interestingly, deficiency for some differentiation drivers increases similarity of 2i profiles to the *in vivo* pre-implantation epiblast. Therefore, we propose that some of the mechanisms driving exit from naïve pluripotency contribute to the lack of complete identity between *in vivo* E4.5 epiblast and ES cells in 2i. This is in line with ESCs representing an engineered interruption in developmental progression and suggests that state-of-the-art self-renewal culture conditions may not yet capture the naïve state perfectly ^1^.

Our analyses provide a comprehensive inventory of genetic factors and regulatory networks governing the major cell fate transition from pre- to post-implantation epiblast (Figure 7B). We present evidence that mammalian development, employs similar regulatory networks operating with similar mechanisms ^61, 62^. These programmed cell state transitions in development do not rely upon instructions delivered by one or two master transcription factors. Rather, a cloud of activity, involving multiple co-ordinated inputs, serves to destabilise an existing, multiply stabilised GRN. Thereby a consolidated cell state can be rapidly dismantled by a combination of signals, triggering transition to the next developmental waystation.

## Acknowledgements

We thank Johanna Stranner, Thomas Sauer and Andy Riddell for help with flow cytometry. Meng Li and Kosuke Yusa for sharing RC9 cells, Andreas Dahl for NGS-sequencing support and Christa Bücker for critical comments on the manuscript. ML is funded by a WWTF-VRG grant (VRG14-006). This study was supported by an FWF/DFG DACH grant to AB and ML (FWF grant number: I 3786; DGF grant number 398882498). AB and MG were supported by the BMBF (Sybacol). RS received support by the Cologne Graduate School of Ageing Research. Wellcome and the Medical Research Council provide core support for the Wellcome-MRC Cambridge Stem Cell Institute. AS is an MRC Professor. This project was initiated within the EU FP7 integrated project SyBoSS (Grant agreement: 242129) co-ordinated by AFS and AB including partners ML and AS.

## Author contributions

AB, ML, AS and AFS conceptualized the study. AB, ML designed the study. AL, ML, AB, RS, MG and FTT designed experiments. AL, MH, JR, PvdL, HFT, LS and MS carried out wet-lab experiments. RS, MG, GGS, FTT and AL performed bioinformatic analyses. ML, AB, AS, AFS wrote the paper with input from AL, RS, MG, MH, FTT and GG. Funding was acquired by ML, AB, AFS and AS.

## Figure Legends

**Supplementary Figure 1.**
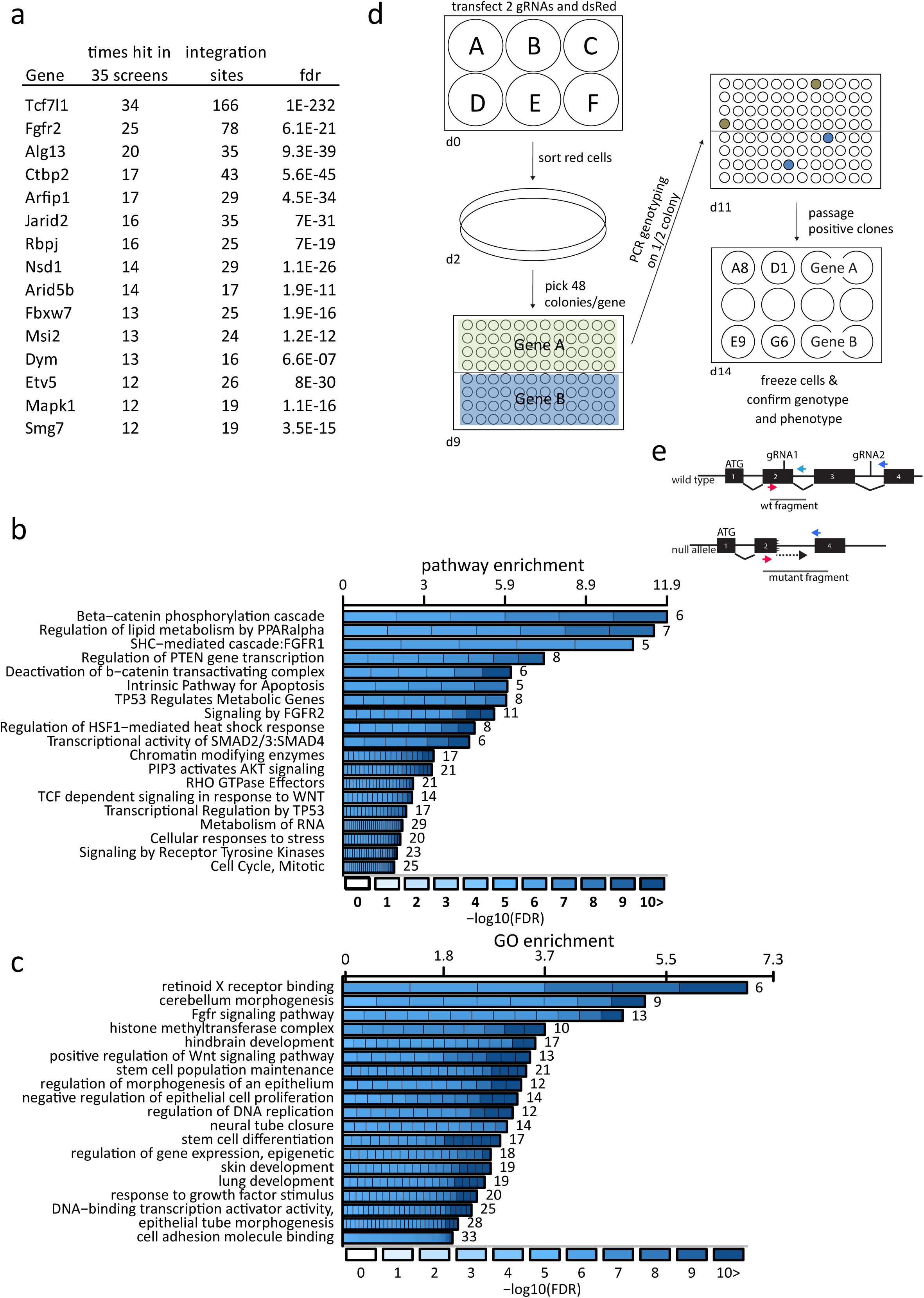
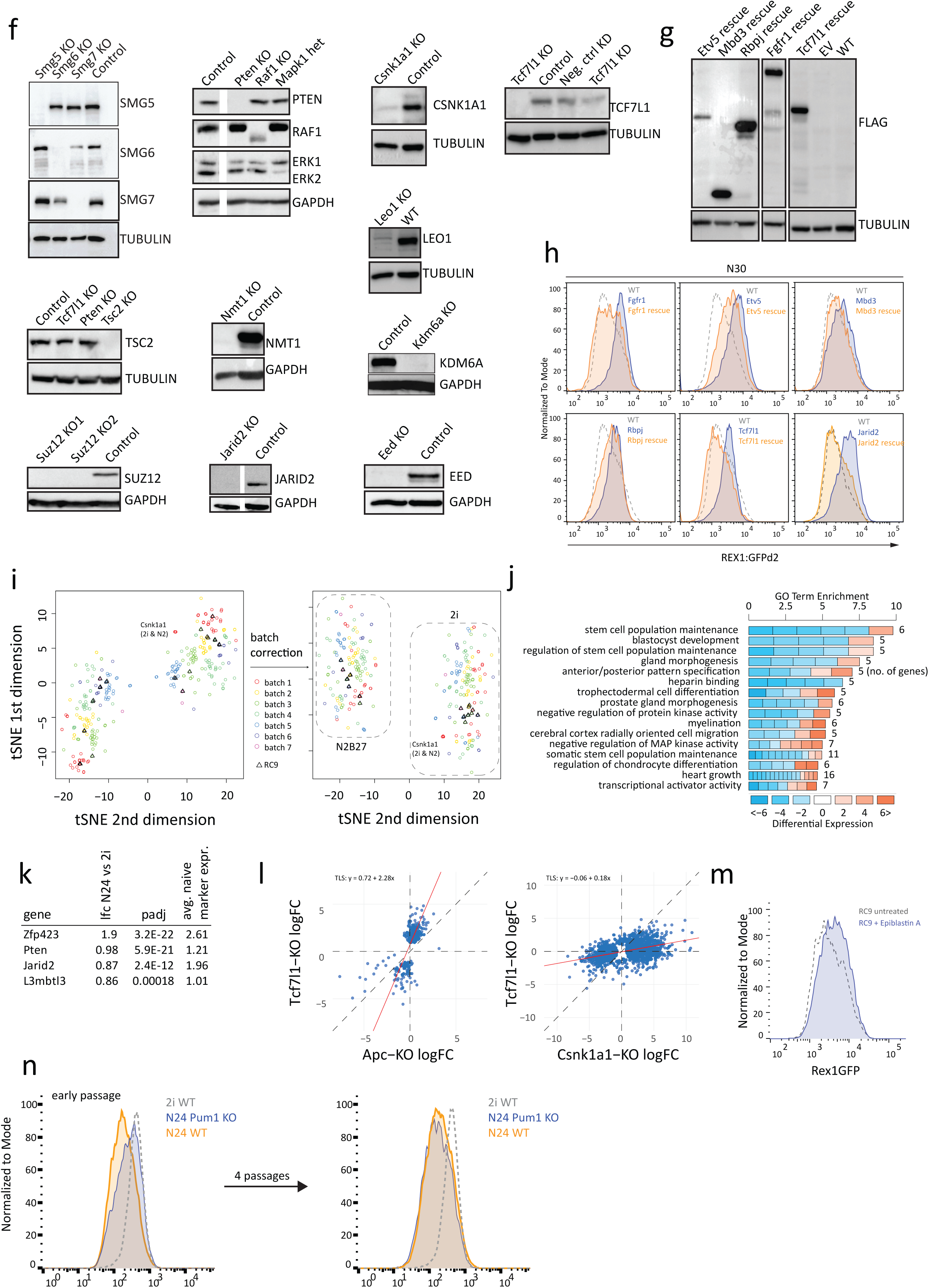
**a,** The top 15 candidate genes from the haploid screen ordered by the number of times the hit was found in the 35 screens, including the number of independent integration sites and the calculated FDR. **b,** Significantly enriched pathways among screen hits ranked according to fold enrichment; colours represent FDR in screen analysis. Numbers next to bars indicate number of hit-genes within category, **c,** Significantly enriched GO terms among screen hits as in **b, d,** Workflow to generate Cas9 knockouts in RC9 ESCs. **e,** Idealized strategy illustrating gRNAs and genotyping primers. **f,** Western analysis using indicated KOs and indicated antibodies. Tubulin or Gapdh were used as loading controls, as indicated. **g,** Anti-flag specific Westerns in indicated KO^rescue^ ESCs upon stable forced expression of 3xflag rescue cDNAs driven from a CAG promoter. **h,** Rex1GFP levels measured by FACS showing restoration of differentiation behaviour in the indicated rescue cell lines at N30. **i,** t-SNE visualization of RNA-seq profiles in 2i and at N24 before and after batch correction. **j,** GO enrichment analysis of the top 10% of genes (by absolute logFC) that were differentially expressed between 2i and N24 in WT ESCs (FDR ≤ 0.05, H0: |FC| < 1.5). **k,** Exit from pluripotency factors showing significant expression changes between 2i and N24 (lfc≥0.8; adj. p-value≤0.05). Log2FC, adj. p value and quantification of the molecular phenotype using the average upregulation of a set of naïve marker genes (*Esrrb, Nanog, Tfcp2l1, Tbx3, Prdm14, Klf4*) are shown. **l,** Dot-plot showing regression analysis of KO-induced changes at N24, comparing *Tcf7l1* KO to Apc and Csnk1a1 KOs. Total least squares (TLS) analysis results are indicated. **m,** Rex1-GFP levels measured by FACS at N24 showing the effect of Epiblastin A addition during differentiation. **n,** Rex1GFP analysis at N24 of *Pum1* KO cells, showing a relatively strong differentiation defect at early passages.

**Supplementary Figure 2.**
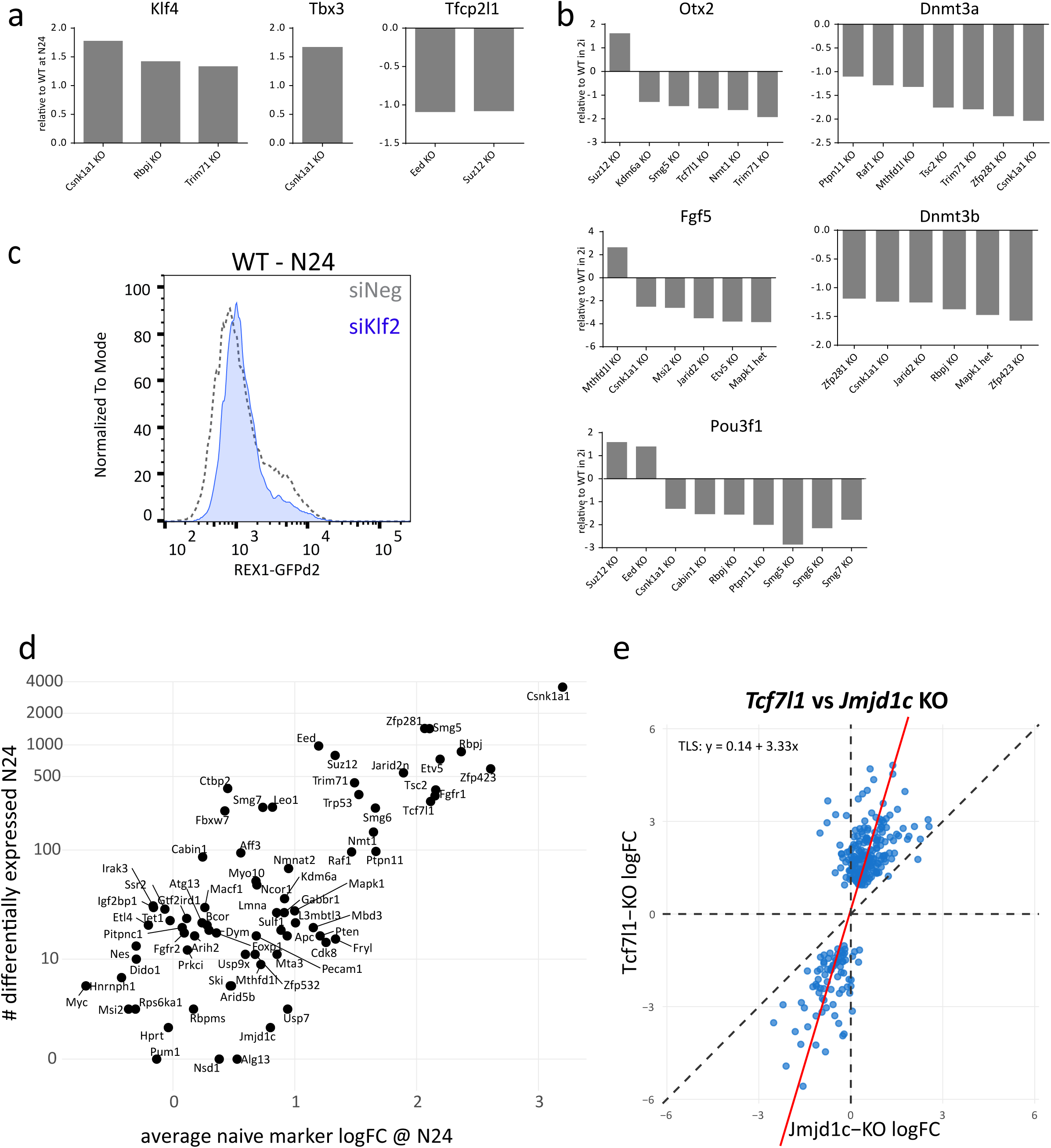
**a,** RNA-seq based fold changes (FC, relative to WT) of indicated naïve marker genes in indicated KOs in 2i (only FCs with adj. p ≤ 0.05 are shown). **b,** RNA-seq derived data showing FCs of indicated formative marker genes in indicated KOs in 2i (only FCs with adj. p ≤ 0.05 are shown). **c,** Rex1-GFP analysis of WT cells at N24, transfected with negative control or *Klf2* specific siRNAs. *Klf2* depletion increases the speed of Rex1GFP downregulation. **d,** Comparison of the number of differentially expressed genes at N24 (FDR ≤ 0.05, H0: |FC| < 1.5) to the mean naïve marker gene expression at N24 (phenotype strength). Stronger differentiation phenotypes correlate with more differentially expressed genes. **e,** Expression log2 fold changes (log2FC) at N24 between knockout and RC9 control, comparing *Tcf7l1* KOs to *Jmjd1c* KOs. Each dot corresponds to one gene. Only genes showing significance (FDR ≤ 0.05, H0: |FC| < 1.5) in either one of the KOs are plotted. Red line: total least square regression; regression coefficients are shown. Convergence on similar target genes is shown by largely identical directionality of regulation of significantly regulated genes in both KOs.

**Supplementary Figure 3.**
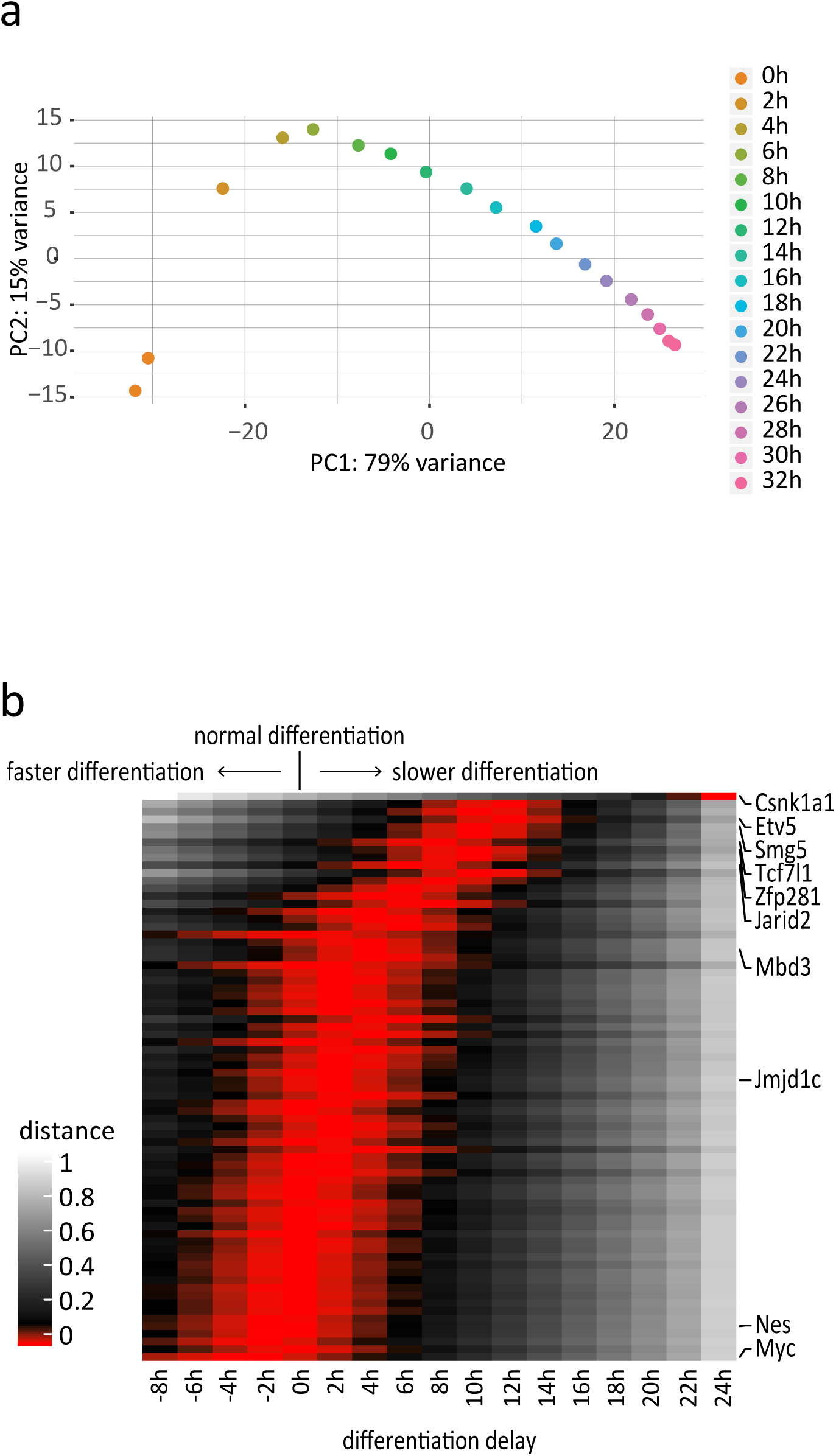
**a,** Principle component analysis (PCA) of the 2h resolved WT differentiation time course sampled by RNA-seq. 0h corresponds to cells in 2i. **b,** Heatmap showing differentiation delays of the 73 KO lines quantified using expression of all 3058 genes differentially expressed in WT between 2i and N24. Red bars indicate closest correlation to that specific time-point. Positive values indicate delayed differentiation. Negative values more rapid differentiation compared to WT. Each line corresponds to one KO. Selected KOs are indicated.

**Supplementary Figure 4.**
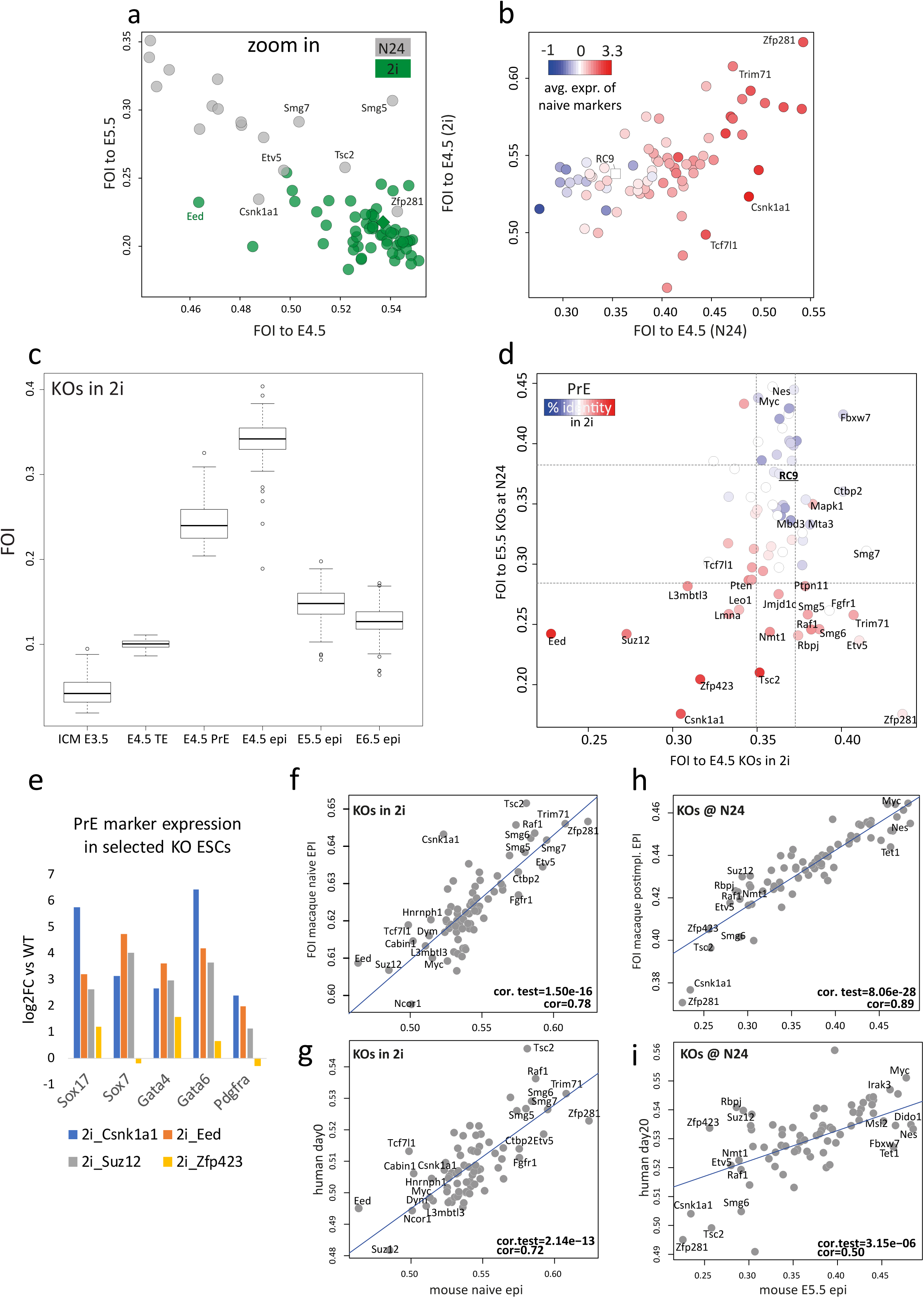
**a,** Zoom in of the plot in Figure 4c, focusing on the overlap of several KOs in 2i (green) and in N24 (grey). N24 samples of *Etv5, Smg7, Tsc2, Smg5* and *Zfp281* KOs show FOIs to the E4.5 EPI similar to KOs cultured in 2i. **b,** Comparison of FOIs to the E4.5 epiblast of all KOs in 2i compared to FOI of all KOs at N24; average naïve marker expression in KOs at N24 is indicated as colour gradient. With some exceptions, stronger *in vitro* phenotypes translate to increased similarity to the E4.5 epiblast in 2i and at N24. **c,** FOIs of KOs in 2i to ICM, E4.5 trophectoderm (TE), E4.5 primitive endoderm (PrE), E4.5 epiblast (Epi), E5.5 Epi and E 6.5 Epi. Analysis extended but similar to Figure 4b. **d,** Plot comparing FOIs of 2i samples to E4.5 epiblast (x-axis) to FOIs of N24 samples to E5.5 epiblast(y-axis). Dotted grey lines indicate 95% confidence intervals. Colour gradient shows similarity of 2i expression profiles to PrE. **e,** Correlation plot of FOIs between 2i profiles and mouse naïve (E4.5) epi and FOIs between 2i profiles and macaque naive epi (correlation significance is indicated). **f,** Correlation plot of FOIs between 2i profiles and mouse naive epi and FOIs between 2i profiles and human naïve ESCs (day0) (correlation significance is indicated). **g,** Correlation plot of FOIs between N24 profiles and mouse E5.5 epi and FOIs between N24 profiles and macaque post-implantation epi (correlation significance is indicated). **h,** Correlation plot of FOIs between N24 profiles and mouse E5.5 epi and FOIs between N24 profiles and human ESCs at day 20 after release into primed culture conditions (correlation significance is indicated).

**Supplementary Figure 5.**
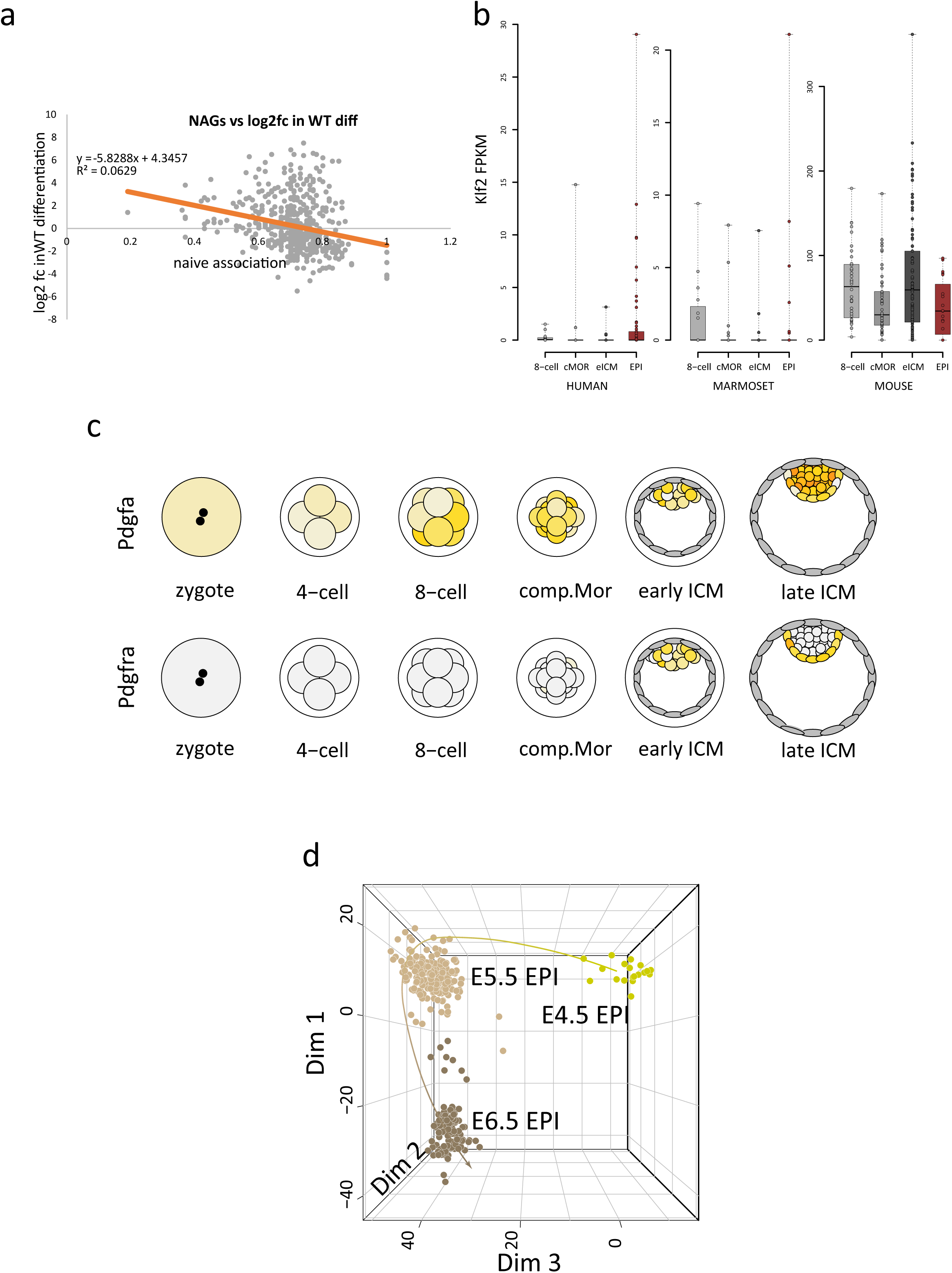
**a,** Expression change of NAGs during naïve to formative differentiation in WT ESCs (y-axis) versus the strength of link to the naïve network (naïve association, x-axis). The orange line indicates the linear regression. The R^2^ value is indicated in the plot. **b,** *Klf2* expression at the indicated stages. Compacted morula (cMOR), expanded ICM (eICM), E4.5 epiblast (EPI) derived from single cell RNA-seq datasets from human, marmoset and mouse preimplantation development. **c,** Plots derived from the GRAPPA visualisation-app showing expression in yellow of *Pdgfa* and its cognate receptor *Pdgfra* during mouse preimplantation embryo development. **d,** Principal component analysis of published single cell RNA-seq datasets showing that Dimension 3 (Dim 3) separates pre- from post-implantation epiblast *in vivo* (as shown in Figure 5a).

**Supplementary Figure 6.**
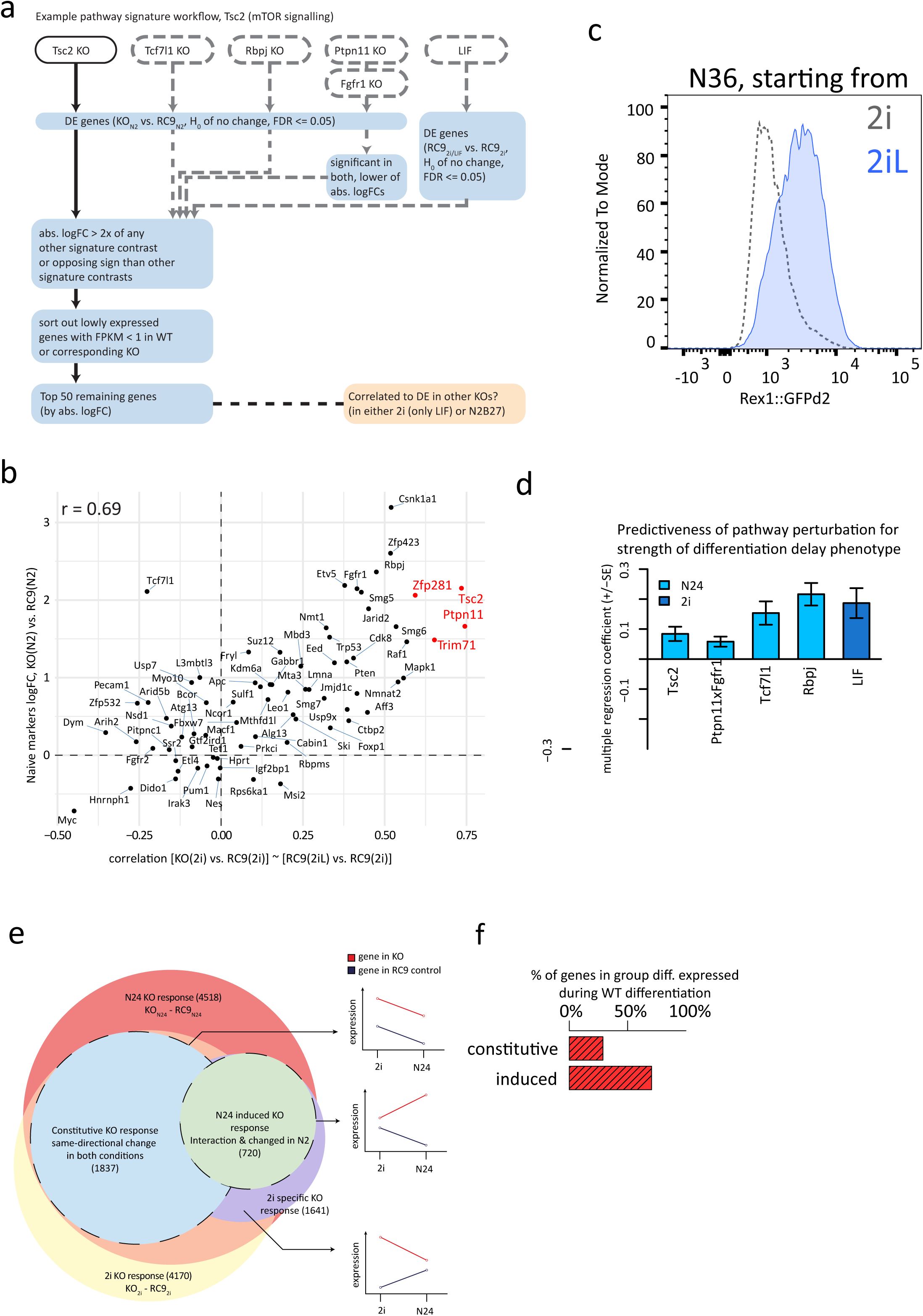
**a,** Schematised computational strategy to derive non-overlapping gene-expression footprints for the five signalling pathways. **b,** Average change in naive marker expression induced by KOs at N24 (y-axis) and their ‘LIF-likeness’ in 2i (x-axis; correlation of RC9 LIF response with knockout induced response in 2i). Overall correlation across these values is r=0.69. The KOs showing the strongest correlations (*Tsc2*, *Ptpn11* and *Trim71* KOs) are indicated in red. **c,** Flow analysis showing delayed Rex1-GFP downregulation in cells starting from cultures in 2i/LIF compared to cells starting from 2i. **d,** Multiple regression coefficients of pathway activity profiles across all KOs when predicting the differentiation delay phenotype (defined by mean naïve marker gene expression change at N24). Error bars indicate the standard error of the coefficient. **e,** Euler diagram showing the overlap of defined gene groups. Panels on the right show exemplary gene expression behaviours. The constitutive and the N24 induced KO response group of genes were used for further analysis. **f,** Percentages of genes in the N24 induced KO response (N24 induced) or constitutive KO response (constitutive) clusters that significantly change during WT differentiation (FDR ≤ 0.05, H0: |FC| < 1.5).

**Supplementary Figure 7.**
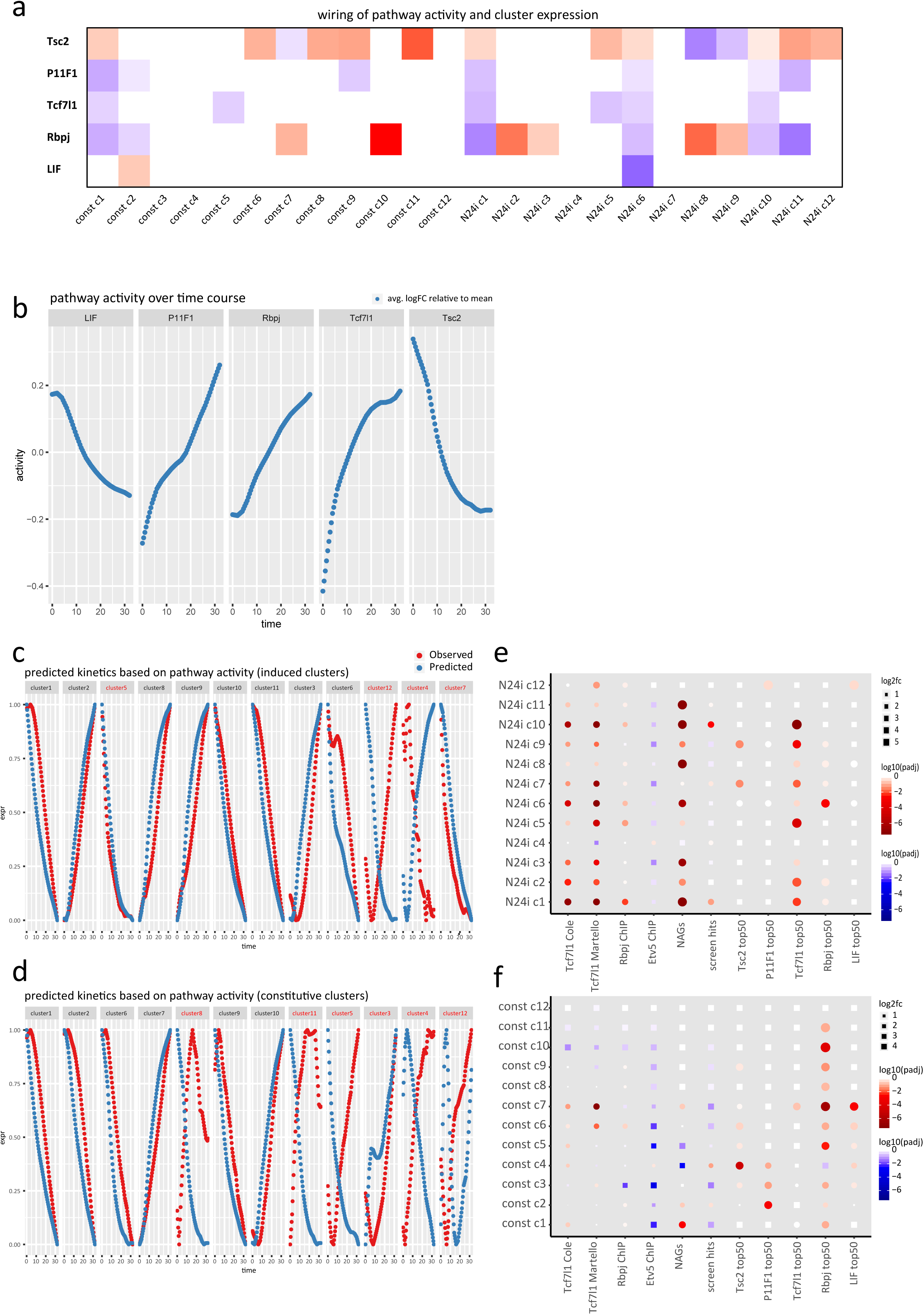
**a,** Heatmap illustrating multiple regression models of pathway activities predicting cluster expression. Only significant (adj. p. ≤ 0.01) interactions are shown. Colour intensity represents the strength of the interaction. Red tiles indicate that higher activity of a certain pathway results in higher expression of the cluster, while blue tiles indicate the opposite. **b,** Pathway activities (average expression of the 50 specific pathway-defining genes) across a WT differentiation time-course for all 5 pathways. y-axis shows average log2FC of pathway defining genes relative to the mean expression throughout the time-course. **c,** Observed cluster expression change (red) and cluster expression change predicted by regression models (blue) throughout the 32h WT differentiation time course for all induced clusters. Predicted cluster expression is calculated using pathway activities over the WT time course and significant (adj. p ≤ 0.01) interactions from the multiple regression models. Graphs show log2FC of cluster expression relative to the mean cluster expression across the time course. Red cluster names indicate that cluster-pathway connections were not predicted in Supplementary Figure 7A or validation through 2h time-course expression failed (predicted expression did not precede observed expression). **d,** Similar to Supplementary Figure 7C for constitutive clusters. **e,** Enrichment from Fisher test of ChIP-targets, NAGs and pathway defining gene sets in induced clusters. Enrichment is indicated by red circles, depletion by blue squares. Size indicates strength of over- or underrepresentation and colour gradient encodes adj. p values. **f,** Similar to Supplementary Figure 7E for constitutive clusters.

## Methods

### Cloning

Guide RNAs (gRNAs) were designed using the computational pipeline described below, or by http://crisprscan.org. To clone the gRNAs for ESC KO of the 154 selected genes, we used RecET recombineering to insert the guide RNA protospacer encoding sequence between the U6 promoter and the universal sgRNA scaffold in pBR322-U6-cm-ccdB-sgRNA-amp, as described previously (Baker et al, 2016). We generated 331 sgRNA expressing plasmids for 149 genes (2 sgRNAs for 131 genes, 3 sgRNAs for 5 genes, 4 sgRNAs for 12 gens and 6 sgRNAs for 1 gene). In brief, the plasmid was linearized at the point of insertion by BstZ17I digestion, purified using the Invitrogen Charge Switch PCR Purification Kit and dissolved in water; 80-mer oligonucleotides containing the protospacer sequence, flanked by 30bp of homology to the target plasmid were dissolved in water. *E. coli* GB05 transformed with the pSC101-Prha-ETgA-tet plasmid ^63^, was cultured to OD600 ∼0.5 in deep-well 96 well plates and then induced for 1 hour for expression of the ETgA operon with L-rhamnose. Electroporation with 200 ng (0.1 pmol) of the linearised vector and 50 pmol of oligonucleotide, using BTX 96 well electroporation plates (MOS96, 2mm gap,) and the BTX ECM630 electroporator with HT-200 adapter, was described previously ^64^. After one hour of recovery time, 100 microliters (1/10th) of the culture was transferred to a new plate with 900 microliters of LB media plus 100 micrograms/ml ampicillin. After overnight selection the saturated cultures were plated for single colonies on L-agar + 100 micrograms/ml ampicillin. A single colony for each construct was picked and the clones were grown under selection in 96 well plates, plasmid DNA was isolated, and Sanger sequenced from both directions using the M13F and M13R primers (flanking the U6-protospacer-sgRNA scaffold cassette, Supplementary Table 6). Sequence analysis confirmed the correct engineering of 316 out 331 plasmids. After sequencing of a second colony or repeat cloning for the remaining 15 clones, all the desired constructs were obtained. Alternatively, annealed oligonucleotides (Supplementary Table 6) were cloned into a BsaI site of a gRNA expression vector (Addgene Plasmid number 41824) ^65^ and correct insertion was determined by Sanger sequencing with the SP6 primer (Supplementary Table 6). For generating rescue cell lines, indicated coding sequences were cloned into a pCAG-3xFLAG-empty-pgk-hph vector ^14^ after PCR amplification and correct insertion was detected by restriction digest and Sanger sequencing with the 3xFlag_seq primer (Supplementary Table 6).

### Cell culture

Diploid (biparental) ES cells were routinely cultured in gelatin coated 25cm^2^ flasks in DMEM supplemented with 15% FCS (batch tested, Biowest), 1x Penicillin-Streptomycin (Sigma), 0.1mM NEEA, 1mM sodium pyruvate, 1mM L-glutamine, 0.05mM 2-mercatpoethanol (Gibco), 10ng/mL LIF (batch tested, in-house), 1.5µM CHIRON and 0.5µM PD0325901 (termed ESDMEM-2i) on gelatin coated plates. The basal medium during all differentiation assays and for haploid ES cell culture (N2B27) consisted of a 1:1 mixture of DMEM/F12 and Neurobasal medium supplemented with 1x B27 (Gibco), 0.5x N2 (homemade), 0.1mM NEAA, 1mM L-glutamine, 1x Penicillin-Streptomycin and 0.05mM 2-mercaptoethanol. Haploid ES cells were routinely cultured in N2B27 supplemented with 3µM CHIR99201, 1µM PD0325901 and 10ng/ml LIF and regularly sorted based on FSC/SSC parameters by FACS ^66^. Haploid Rex1GFPd2-IRES-BSD (hRex1GFPd2, ^67^ and haploid Rex1::mKO2-IRES-BSD/pOct4-GFP-IRES-Puro ESCs (generated in this study from a double reporter mouse) were derived from activated oocytes in 2i/LIF medium and used for screens. Biparental diploid ES cells with a GFP driven by the endogenous Rex1 promoter and EF1a driven Cas9 targeted to the Rosa26 locus (RC9 cells, ^13^ served as parental cell line for the KO lines generated in this study. A list of KO cell lines can be found in Supplementary Table 6.

### Transposon-based saturation screen for key players of the exit from pluripotency

The genetrap vectors 5’-PTK-3’, TNN, TNP (Horie et al. 2011, Allan Bradley Lab) together with haploid Rex1::GFPd2 ES cells (hRex1GFPd2, Leeb et al. 2014) and haploid Rex1::mKO2-IRES-BSD/pOct4-GFP-IRES-Puro ES cells (hOR, this study) were used in 35 independent experiments to drive the screen to saturation. Mutant pools were generated by electroporation of 10^7^ haploid ESCs using a GenePulser Xcell (270 V, 500 μF, ∞ Ω, Biorad) with 0.5 μg genetrap plasmid and 10 μg hyperactive transposase (hyPBase) ^68^. 24h after electroporation, selection was started for 4 days using 1µg/ml Puromycin. Thereafter cells were plated at a density of 10^4^ cells/cm^2^ in N2B27 medium in the absence of LIF or inhibitors to allow differentiation. After 7–10 days in differentiation conditions, GFP-positive cells were sorted and replated at a density of 10^4^ cells/cm^2^. After culture in N2B27 medium for a further 7–10 days, GFP-positive ESCs were resorted, expanded for 48h in 2i/LIF medium and DNA was isolated using the PureGene kit (Qiagen) according to the manufacturer instructions. Sequencing libraries were prepared using an optimized Splinkerette PCR protocol (Leeb et al. 2014) with a modified and improved set of adapters and primers (see Supplementary Table 6). Adapters were annealed at 50µM each in T4 DNA ligase buffer by incubating at 97.5°C for 150 sec followed by a temperature decrease of 0.1°C/5sec for 775 cycles. Ready-to-use adapters were stored at - 20°C. Genomic DNA was isolated using the PureGene kit (Qiagen) according to the manufacturers recommendations and quantified on the Qubit 2 fluorometer (Life Technologies) using the Qubit dsDNA Broad Range Assay (Life Technologies, Q32853). 2µg of genomic DNA was diluted to a total volume of 120µl with Low TE buffer (10mM Tris-HCl, pH 8.0, 0.1mM EDTA) and sheared to 250bp on a Covaris S-series Sample Preparation System with the following settings: Duty Cycle – 20%, Intensity – 5, Cycles per Burst – 200, Time – 60 seconds, Temperature – 4° to 7°C. After shearing DNA was isolated with the QIAquick PCR purification kit (Qiagen) and quality assessed using the Agilent Bioanalyzer with the DNA high sensitivity kit (Agilent). End-repair was performed using the NEBNext End Repair Module (E6050S) by mixing 100µl sheared DNAl, 11.6µl buffer and 6µl enzyme followed by incubation at 20°C for 60min. After cleanup with the PureLink PCR Purification Kit (K3100-01), 42µl of the end repaired DNA was A-tailed with the NEBNext dA-Tailing Module (E6053S) with 5µl buffer and 3µl enzyme at 37°C for 60 minutes. Following PureLink purification of to the A-tailed DNA, the annealed adapters were ligated using theNEBNext Quick Ligation (E6056S) kit at 20°C for 30min. Products were cleaned up using Ampure XP beads at a 1:1 ratio. Libraries were PCR amplified using Kapa HIFI (Roche) with 10µM PB5_pr1 and 10µM SplAP1 during 18 cycles of 20sec 98°C, 20sec 63°C and 40sec 72°C with an initial denaturation at 95°C for 2minutes and a final extension at 72°C for 5min. Samples were cleaned up using Ampure XP beads at a ratio of 0.8. A second round of PCR was conducted using xxul from PCR1 using the Kapa HIFI polymerase with 10µM SplAP2 and 10µM index primer for 13 cycles at an annealing temperature of 60°C with all other settings as in PCR1. Final libraries were quantified using the KAPA SYBR Fast qPCR Mix with primers SYB7 and SYB5 at 10µM each and a final quality control was conducted with the Agilent Bioanalyzer DNA High Sensitivity Assay.I ntegration sites were mapped as in ^12^ and described below.

### Generation of KO ESCs

Direct comparison between KOs necessitates a similar culture and passaging history. We therefore established an experimental pipeline to parallelise the generation of multiple KO ESC lines per week. 2×10^5^ RC9 cells were transfected in a 6-well format with a set of two gRNA containing vectors (Supplementary Table 6; (1µg each), together with 0.5µg of pCAG-dsRed by Lipofectamine2000 (3µl per reaction). 12-16h after transfection, medium was replaced. To enrich for transfectants, cultures were sorted for GFP/DsRed double positive cells on a BD FACS Aria III 48h after transfection and plated at clonal density in 6cm dishes (100 to 800 cells per dish) in ESDMEM2i. Approximately one week after sorting, 48 colonies per gene KO were picked into 96-well plates, trypsinized and split into one expansion-plate and one plate for PCR genotyping after boiling lysis and proteinase K treatment (see PCR genotyping). Identified KO clones were expanded to 12-well or 6-well plates after 3 days of growth and the remaining plate was frozen in 50%FCS/10% DMSO as a backup. Clones were frozen from 12- or 6-well plates in freezing medium containing 50% FCS and 10% DMSO. Of the attempted 154 gene disruptions, we achieved 115 homozygous knockout cell lines. In 16 cases, only heterozygous clones were obtained (indicating essentiality of these genes), and in 23 cases we were unable to recover gene disruptions, despite multiple attempts (Supplementary Table 6).

### PCR genotyping

Crude DNA lysates were generated by pelleting half of a picked colony in a PCR plate at 500g at 4°C, followed by two washes with PBS. 25µl of water were added and pellets were boiled for 5min at 95°C. After cooling, Proteinase K was added at a final concentration of 3µg/µl and incubated at 65°C for 1h followed by inactivation at 95°C for 10min. Successful KO generation was confirmed by PCR, employing a three-primer strategy. Specific reverse primers for the deletion event and a possible wildtype allele were used in combination with a common forward primer (Supplementary Table 6). Cell lines with indels caused by a NHEJ event around the exonic gRNA1 target site, were detected by Sanger sequencing of the “wildtype”-band. Verification of the KO-events was performed by the same PCR strategy on DNA, isolated using the PureGene DNA isolation kit (Qiagen) according to the manufacturer recommendations. For genotyping PCR, we used OneTaq (NEB) or JumpStart RedTaq (Sigma) PCR Master Mix according to the manufacturers recommendations for cycling and a standard annealing temperature of 55°C and 35-40 cycles.

### Immunoblotting

Whole cell fractions were isolated using RIPA buffer and protein concentrations were determined using a Bradford Assay (Bio-Rad). 20µg whole cell lysate were separated on 8-12% SDS-PAGE gels (depending on the molecular weight of the target proteins), and subsequently blotted on 0.2µm nitrocellulose membranes (Amersham). Membranes were blocked in TBS or PBS containing 5% milk and 0.1% Tween-20 or 3% BSA and 0.1% Tween 20 for 1h. Primary antibodies were incubated overnight at 4°C, at a dilution of 1/1000 with the following exceptions: Anti-Csnk1a1 1/200, Anti-Flag M2 1/1000 Anti-Gapdh 1/25000, Anti-Tubulin 1/5000, Anti-Smg5 1/200, Smg7 1/2000. Secondary goat anti-mouse IgG HRP (1:10000) or goat anti-rabbit IgG HRP (1:15000) were incubated at RT for 1h in blocking solution. Antibody binding was detected using the ECL select detection kit (Amersham). A list of antibodies can be found in Supplementary Table 6.

### Parallel Differentiation

ES Cells were differentiated in seven batches, always including WT controls, control KO ESCs (no phenotype expected) and a range of ESCs with expected weak to strong phenotypes. Where applicable two independent KO clones were used, other KO ESCs were cultured in parallel replicates before intiation of differentiation. 24h before starting the differentiation assay, medium was changed from ESDMEM2i to N2B27 based 2i/LIF medium. Cells were then trypsinized, counted and plated in 2i (without LIF) at a density of 10^4^ cells/cm^2^ in two replicate wells of a 6-well plate for differentiation, one 6-well as undifferentiated control and 2 wells of a 12-well plate (undifferentiated control and differentiated sample) to monitor differentiation by FACS. 12h after plating, cells were washed twice with PBS and medium was changed to unsupplemented N2B27. For the undifferentiated controls medium was replaced with fresh 2i. 24h later, at N24, cultures were harvested in RLT buffer and stored at −80°C before isolation of RNA using the RNeasy Mini Kit (Qiagen). mRNA was isolated from 1 ug total RNA by poly-dT enrichment using the NEBNext Poly**a,** mRNA Magnetic Isolation Module according to the manufacturer’s instructions. Samples were eluted in 15ul 2x first strand cDNA synthesis buffer (NEBnext, NEB). After chemical fragmentation by incubating for 15 min at 94°C the sample was directly subjected to the workflow for strand specific RNA-Seq library preparation (Ultra Directional RNA Library Prep, NEB). For ligation custom adapters were (Supplementary Table 6). After ligation, adapters were depleted by an XP bead purification (Beckman Coulter) adding beads in a ratio of 1:1, followed by an index PCR (15 cycles) using Illumina compatible index primer. After double XP beads purifications (with beads added in a ratio 1:1) libraries were quantified on a Fragment Analyzer run with a NGS Assay Kit (Agilent) and loaded on a HiSeq 3000 flowcell with 50 cycles single end sequencing by pooling the samples based on molarity aiming for 30 mio. reads per sample.

### 2h resolved differentiation timecourse

Medium was changed to N2B27-2iLIF 24h before the assay. Cells were trypsinized, counted and plated at a density of 10^4^ cells/cm^2^ in 6-well plates in 2i (without LIF). 12h later medium was changed to unsupplemented N2B27 (or 2i for the undifferentiated controls) and cells were harvested in buffer RLT every 2 hours for the next 32h (N32). RNA was isolated using the RNeasy Mini Kit (Qiagen). All libraries were generated from 500ng RNA using the TruSeq Stranded mRNA Library Prep Kit (Illumina) and analysed on a HiSeq4000, Paired End 75bp.

### LIF signature

1×10^4c^ells/cm^2^ were plated in N2B27-2i and N2B27-2iLIF and grown for 24h after which RNA was isolated using the RNeasy Mini Kit (Qiagen). Libraries were made from 1µg high quality RNA using the Quantseq 3’ mRNA-Seq Library Prep Kit FWD for Illumina (Lexogen) according to the manufacturer’s instructions. Libraries were quantified using qPCR according to the Quantseq Library Prep Kit and with the included primers, pooled and quality checked using an Agilent Bioanalyzer DNA High Sensitivity Assay (Agilent). Multiplexes were sequenced on a HiSeq4000, Single End 50bp.

### RNAi experiments

For RNAi FlexiTube siRNAs against Klf2 were used with AllStars Negative Control siRNAs (Qiagen). 20ng siRNAs/4×10^4^ cells were transfected in 2i using DharmaFect (Dharmacon). 12h later medium was changed to N2B27 after two PBS washes. At N24 and N32 Rex1GFPd2 expression was determined by flowcytometry (see Flowcytometry Analysis) and cells were harvested in RLT buffer and isolated with the RNeasy mini kit (Qiagen) according to the manufacturer’s recommendations. cDNA was transcribed using the SensiFAST cDNA Synthesis Kit (Bioline). Naïve and primed marker expression was determined by qPCR using the Sensifast SYBR No Rox-Kit (Bioline). Primers are listed in Supplementary Table 6.

### Flow cytometry

Cells were dissociated in 0.25% trypsin/EDTA and trypsin was neutralized with DMEM supplemented with 15% FCS. After passing through a 40µm mesh, Rex1-reporter activity was measured using a BD Fortessa machine. High-throughput-measurements were performed in 96-well plates using the HTS unit of the BD Fortessa. Data were analyzed using FlowJo software.

### Data Analysis

#### Transposon Mutagenesis Screen in Haploid Murine Embryonic Stem Cells

Integration mapping was performed as previously described ^12^. Most gene-trap insertions were found in intronic sequences, consistent with the relative length distributions of genomic features. However, after normalising to available TTAA sites per region, TTAA sites in 5’UTRs and promoters (<-500bp) were overrepresented, whereas TTAA sites in introns were relatively depleted. 7760 independent integrations passing cut-off criteria were mapped to 3469 genes. Of those, 232 Genes (2474 independent integration sites) were hit in five or more independent screens, indicating that those are integrations causative for the detected differentiation defect. To complement the candidate list and correct for gene specific biases we further developed a strategy to assign statistical strength to candidate genes by assessing the number of integrations per gene in relation to available TTAA transposon integration sites We utilized the number of different TTAA-sites within each gene to calculate the probability of having integrations in *k* or more different locations by chance. All TTAA sites in mm10 were assessed using bowtie and the analysis was restricted to genes classified as protein coding, long non-coding RNAs and micro RNAs. We computed the probability that a gene is hit *k* times given that it contains *n* TTAA transposon integration sites assuming a binomial distribution (while *k* is the observed number of integrations).

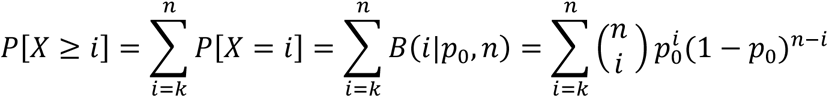

*p_0_* was then calculated by 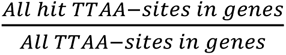

We applied this test to genes with at least 26 TTAA sites, as genes with fewer TTAA sites would be significant (p-value < 0.05) with a single integration. Candidate gene lists between both analyses showed overlapping results and top hits were largely identical. This resulted in a list of 421 significant genes (adjusted p-value < 0.01 and hit in at least two independent screens).

#### Guide RNA design criteria

To automate the design of sgRNAs as far as possible we used the R package CRISPRseek (Zhu et al., 2014) to score the efficiency and likely off-target binding of candidate sgRNA sequences. For a selected gene the algorithm proceeds as follows:

1. Retrieve the sequence of the first protein coding exon
2. Sequence has to be at least 50 bases long
3. Find sgRNAs within the sequence and score them.
4. If no appropriate sgRNA could be found, move to next exon
5. Still in the first third of the coding sequence?
6. Find a second sgRNA in an intron at least 5kb away, but no more than 30kb
7. Score the sgRNAs

A minimal efficiency of 0.2 and an off-target score < 100 was required.

#### PCR Primer Design Strategy for Simplified Knock-out Validation

The surrounding regions of the sgRNAs (min distance 20bp) were used as input for PRIMER3 primer predictions. Primers were designed to have a length of 18-27 bases with an optimum of 20 and to have melting temperatures between 57°C and 63°C with an optimum of 60°C. All primers were checked for off-target binding sites by using Blastn. Blastn was called with the parameters:

-task megablast -use_index false -word_size 7 -perc_identity 65 –evalue 30000 - max_target_seqs 50 -num_threads 4 –outfmt 7.

#### RNA quantification, RNAseq

Quality control was performed using fastQC (version 0.11.5). Transcripts were mapped to the mm10 mouse reference genome and counts determined using STAR (version 2.5.3). RPKM values were calculated using DeSEQ2 (version 1.20.0).

#### RNA quantification, QuantSeq

Transcripts from RC9 cells in 2i and 2i/LIF medium were measured using QuantSeq (Lexogen) and quantified as described. Reads were trimmed with bbduk (version 35.92). Quality control was performed using fastQC (version 0.11.5). Transcripts were mapped to the mm10 mouse reference genome using STAR (version 2.5.3). After indexing with samtools (version 1.3) reads in genes were counted using HTSeq-count (version 0.6.0). Counts were subjected to differential expression analysis.

#### Differential expression analysis

Differential expression analysis was carried out using limma (version 3.30.13), after transforming transcript counts using the included voom function. We then fitted a linear model to sample measurements from each combination of knockouts (including RC9 control) and conditions (2i and N24). RNAseq batches were included in the model as a confounding factor. Contrasts were fitted to determine the expression differences between a) each knockout sample in 2i and the RC9 control, b) each knockout sample in N24 and the RC9 control, and c) the difference between the log-fold-changes calculated in a) and b) (interaction effect). We also fitted a contrast to determine changes during normal differentiation, i.e. the differential expression between RC9N24 *versus* RC92i. All fitted contrasts were tested for differential expression using moderated t-statistics through the limma functions eBayes (H0 of no change) or treat (H0 interval [-log2(1.5), log2(1.5)]). p-Values were corrected for multiple testing using the Benjamini Hochberg method (FDR). For subsequent analyses the more stringent p-values and FDR values as calculated by treat were used (unless noted otherwise). Differential expression analysis of 2i/LIF vs. 2i RC9 measurements was carried out separately, but in an identical fashion.

#### GO enrichment analysis

Mouse gene GO annotations were extracted from the R package org.Mm.eg.db (version 3.4.0). GO terms that included between 5 and 500 genes were selected for further analysis. For each gene list of interest, significance of functional enrichment of GO terms compared to the background list was determined using Fisher’s exact tests. The background was restricted to genes whose transcripts were detected at a median count above 5. To reduce redundancy of functional categories, we clustered GO terms that differed in 5 or fewer genes of interest (using the R base function hclust on the L1-distance of the binary membership matrix). The smallest GO term by total annotations was selected independently from Fisher test results as the primary, i.e. most specific term to represent each cluster. We then carried out correction for multiple hypothesis testing using the Benjamini-Hochberg method on all primary GO terms to obtain adjusted p-values.

#### Naive marker-dependency of N24 KO expression patterns (NAGs)

To quantify the similarity of each gene’s differential expression (DE) pattern across KOs at N24 with the DE pattern of seven core pluripotency markers (Nanog, Esrrb, Tbx3, Tfcp2l1, Klf4, Prdm14 and Zfp42), multiple regression analysis was performed. I.e., each gene’s DE pattern was modelled as a linear combination of marker DE patterns, quantifying the strength of association as their shared variance (multiple-R^2^). NAGs were defined as having a naïve marker R^2^ of ≥ 0.7.

#### Cluster analysis (constitutive & N24 induced)

Based on the responses to the knockout conditions (excluding *Suz12*, *Eed* & *Csnk1a1* KOs) in both N24 and 2i and their interactions with the RC9 control, we defined two main categories of responsive genes: (1) the constitutive knockout response: genes that were significantly changed (adj. p-≤ 0.05) in the same direction in both N24 and 2i, in at least one knockout condition. Genes from the N24 induced KO response category were excluded. (2) N24 induced knockout response: genes that were significantly changed in both N24 and the knockout:RC9 interaction term, in at least one knockout condition. To determine whether genes in the N24 induced knockout response subset were overall more correlated with naive pluripotency markers, we checked the distribution of naive marker multiple-R2 values in the ‘N24 induced knockout response’ set compared to both the background distribution and genes classified as part of the constitutive KO response.

To determine if distinct functional clusters of genes that may be activated in all, single, or sub-groups of knockouts, we clustered genes in either of the categories (constitutive and N24 induced response) based on their KO^N2^ vs. RC9^N24^ log-fold-changes, using the R base package hclust with the “Ward.D2” method. We determined the total within-cluster variance at different numbers of clusters. The number of clusters was chosen such that the model improved strongly by increasing the number of clusters up to that value, but only marginal gains result from increasing it further. Thereby 12 gene clusters in the constitutive response and 12 clusters in the induced response were identified. GO enrichment analysis was carried out as described above for member genes of each cluster

#### Differentiation pathway correlation analysis

We defined downstream responses of key differentiation pathways using the most specific responses to the knockouts of their upstream regulators: *Tsc2* (mTOR), *Ptpn11* & *Fgfr1* (Fgf/Erk signalling), *Tcf7l1* (Wnt signalling), and *Rbpj* (Notch signalling). In addition, LIF-specific changes were defined as the response of genes in RC9 cells in the 2i/LIF compared to the 2i condition. To ensure specificity, we required that the log-fold-change of each response gene for a given regulator (in N2B27 medium) exceeded that observed in the remaining conditions by a factor of two. In the case of Ptpn11 & Fgfr1, this criterion had to be met by the lower of the two log-fold-changes for each gene. Additionally, the genes were required to satisfy a FPKM (CPM in case of LIF) cut-off of at least 1 in either the KO or the WT. From all remaining, significantly expressed genes (FDR ≤ 0.05) we then selected, by absolute log-fold-change in descending order, the top 50 most specific genes for each pathway (Supplementary Figure 6a).

Pathway activity was determined across all other knockouts at N24. For each pathway, the log-fold-changes of the 50 pathway-specific response genes from the associated regulator knockout (KO^N24^ vs. RC9^N24^) were selected. We then correlated these log-fold-changes to the log-fold-changes of the same genes in all other knockouts. A positive correlation indicates that the pathway is disrupted in a manner similar to knocking out its upstream regulator, whereas a negative correlation indicates the opposite. In the case of LIF-specific genes, a positive correlation indicates an effect similar to that of adding LIF. The analysis was repeated for all 2i experiments (using the same lists of specific genes as in N24). P-values of correlation values were calculated based on Fisher’s Z transform. P-values were then corrected for multiplicity using the Bonferroni method.

To determine how predictive the calculated correlations were of the differentiation phenotype (mean naive marker logFC vs. RC9 in N2B27), we carried out multiple linear regression and extracted the coefficients of the model to establish the relative contribution of each pathway signature to naïve marker deregulation.

#### Quantification of differentiation delay in knockouts

Raw RNA sequencing data was aligned to the mm10 genome and read counts were obtained using the STAR aligner (version2.5.2b). Gene expression for the following biotypes was quantified: protein_coding, misc_RNA, miRNA, scaRNA, scRNA, snoRNA, snRNA, sRNA. Read counts were normalized using DESeq2 (version 1.22.2) and subsequently the log2fc between all samples and the mean expression of the 2i samples was calculated. This resulted in expression profiles for each gene over 32 hours of differentiation.

Gaussian process regression was used to smoothen the profiles of each gene (R package tgp version 2.4-14). The expression profiles of 73 knockouts at N24 were mapped on the differentiation axis from the time course to quantify differentiation delays per KO, i.e. we aimed to quantify how many hours the KO expression pattern is ‘behind’ the expected differentiation in WT cells. This ‘mapping’ to the time axis was done first computing the logFCs between N24 versus 2i for each KO. The resulting logFC profiles were compared to the logFCs at each time point during the WT differentiation. We assumed that the time point at which the difference between the WT profile and the KO profile is minimized best reflects the molecular differentiation state of the respective KO.

We used the Euclidean distance for this purpose. To make distances better interpretable between the knockouts the distances were set to:

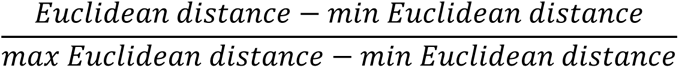

where the maximum and minimum distances refer to the respective maximum and minimum Euclidean distances of the respective KO across all time points (i.e. the worst and the best matching time point). Thus, the best matching time point will get a distance score of 0.

In order to further increase the time resolution, additional time points were imputed via linear interpolation. Here we interpolated expression profiles every 15 minutes between the two neighbouring time points of the time point with the smallest Euclidean distance. Timing of knockouts was repeated on the interpolated time points and the new minimal distance was used to quantify the differentiation delay.

#### Prediction of cluster expression by pathway activity

Pathway activities for each pathway in all KOs were calculated using following formulas:

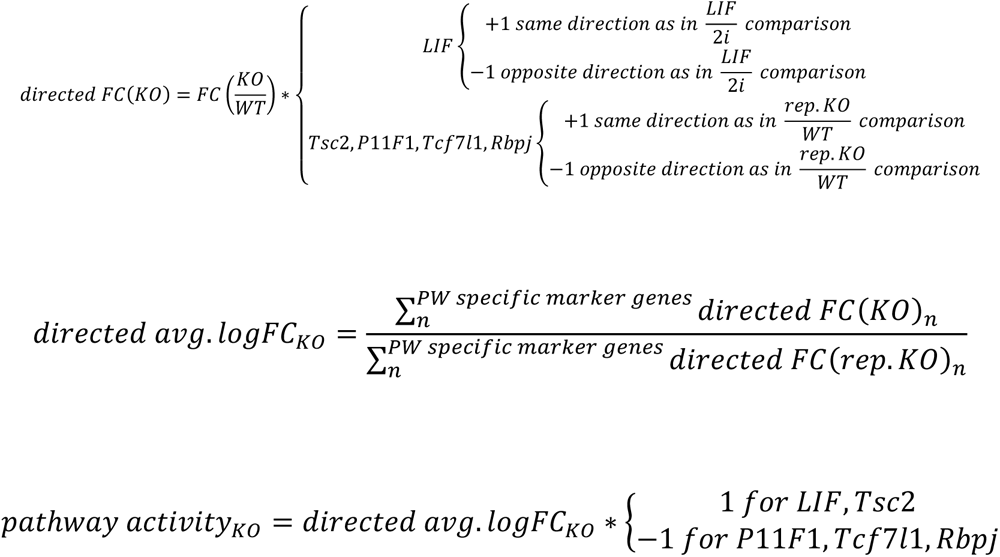

The combinations of pathway activities and average cluster expression changes over all KOs were used to build a multiple regression model to predict the expression changes in each cluster by the five pathway activities. Connections between pathway activities and cluster expression changes not making a significance cut-off (padj <= 0.01) of the multiple regression models were excluded from each model. The remaining model was tested in the time course data. Connections from clusters where prediction of expression changes based on pathway activities did follow or contradict the observed expression changes in the time course were excluded.

#### Embryo comparison

Sequencing data were obtained from the European Nucleotide Archive ^69^ from single-cell mouse embryo profiling studies ^10, 70^. Human capacitation dataset was downloaded from Gene Expression Omnibus website (GSE123055). The macaque FKPM expression dataset was provided by ^33^. Orthologues genes mapping in 1-to-1 fashion were used during comparative analysis. Alignments to gene loci were quantified with htseq-count ^71^ based on annotation from Ensembl 87 ^69^. Principal component and cluster analysis were based on log2 FPKM values computed with the Bioconductor package DESeq ^72^, custom scripts and FactoRmineR packages ^73^. High variable genes were calculated by fitting a non-linear regression curve between average log2 FPKM and the square of coefficient of variation. Specific thresholds were applied along the x-axis (log2FPKM) and y-axis (coefficient of variation) to identify the most variable genes. Fractional identity between the bulk RNAseq (2i, N24) and the mouse, macaque embryo or human capacitation dataset were computed using R package DeconRNASeq ^74^.

#### GENEES (<u>Ge</u>netic <u>N</u>etwork <u>E</u>xplorer for <u>E</u>mbryonic <u>S</u>tem Cell Differentiation)

To both visualize and interrogate gene expression analysis of all KO ES cells we have set up the online tool GENEES (<u>Ge</u>netic <u>N</u>etwork Exploration of Embryonic Stem Cell Differentiation). It is accessible at [http://shiny.cecad.uni-koeln.de:3838/stemcell_network/; User: stemcells; Pw: S3m7KmIfHr9]. The app integrates multiple visualisations and summary statistics to make all information, from the gene level to network modules / pathways / GO terms, directly available for exploratory analysis. The app has 4 different tabs for inspection of single genes, predefined genesets, pre-computed clusters and custom genesets. Single genes can be inspected in the “Genes tab”, while predefined genesets are available for selection under the “Genesets” tab. The previously described constitutive clusters and InteractionN2 clusters can be further inspected in the “Pre-computed clusters” tab. In the “Select custom geneset” tab, a set of genes of interest subjected to analysis.

